# Peristaltic pump-triggered amyloid formation suggests shear stresses are *in vivo* risk factors of amyloid nucleation

**DOI:** 10.1101/2024.09.16.613380

**Authors:** Yuji Goto, Tomoki Ota, Wenlou Yuan, Ikuko Yumen, Keiichi Yamaguchi, Hirokazu Matsuda, Suguru Yamamoto, Hirotsugu Ogi

**Affiliations:** Graduate School of Engineering, Osaka University, Osaka, Japan; Graduate School of Medical and Dental Sciences, Niigata University, Niigata 951-8510, Japan

**Keywords:** α-synuclein, amyloid β, amyloid fibril, β2-microglobulin, hen egg white lysozyme, peristaltic pump, shear stress, supersaturation

## Abstract

Amyloid fibrils, crystal-like fibrillar aggregates of denatured proteins, are formed linked with the breakdown of supersaturation, causing a series of amyloidosis including Alzheimer’s and Parkinson’s diseases. Although varying *in vitro* factors are known, *in vivo* factors breaking supersaturation are unclear. We found that flowing by a peristaltic pump effectively triggers amyloid formation of hen egg white lysozyme, a model amyloidogenic protein, and, moreover, amyloidosis-associated proteins (i.e. α-synuclein, amyloid β 1-40, and β2-microglobulin). The peristaltic pump-dependent amyloid formation was visualized by a fluorescence microscope with looped flow system, revealing dynamic motions under flow. Among them, amyloid fibrils of amyloid β 1-40 were stickier than others, self-associating, absorbing to loop surfaces, and surging upon flicking the loop, implying early stages of cerebral amyloid angiopathy. On the other hand, β2-microglobulin at a neutral pH showed unique two-step amyloid formation with an oligomeric trapped intermediate, which might mimic amyloid formation in patients. Peristalsis-caused strong shear stresses were considered to mechanically break supersaturation. Shearing stresses occu*r in vivo* at varying levels, suggesting that they break otherwise persistent supersaturation, thus triggering amyloid formation and ultimately leading to amyloidosis. (182 words <200 words)

## Introduction

Amyloid fibrils, crystal-like fibrillar aggregates of denatured proteins^1–6^ associated with a series of amyloidosis including Alzheimer’s and Parkinson’s diseases^7,8^, are formed linked with the breakdown of supersaturation^9,10^. Amyloid fibrils have been reproduced *in vitro* even in the absence of seeds under a variety of conditions by distinct mechanisms^9,10^. These include: (i) a counter ion-binding mechanism observed under acidic conditions in the presence of moderate concentrations of salts, (ii) a salting-out mechanism observed under high salt conditions independent of pH, (iii) a hydrophobic additive-binding mechanism observed in the presence of moderate concentrations of alcohols, detergents like sodium dodecyl sulfate (SDS), or membrane surfaces, and (iv) pI-precipitation under low salt conditions. Common to these mechanisms is the establishment of a supersaturated state of responsible proteins and subsequent breakdown^10–15^.

Analogous to crystallization, spontaneous amyloid formation occurs in proportion to the degree of supersaturation (*S*)^16^:

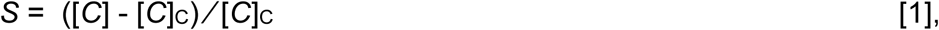

where [*C*] and [*C*]_C_ are the initial solute concentration and thermodynamic solubility (i.e., critical concentration), respectively. Varying conditions as described above increase *S* and, thus, the risk of amyloid formation^10,17^. However, under a metastable condition where supersaturation persists in the absence of seeding, mechanical triggers are required to break supersaturation^18^. Thus, although stirring or ultrasonication is often employed as a mechanical trigger of amyloid formation, we do not know triggers *in vivo*.

Fluid flow stresses have been increasingly focused on as mechanical triggers initiating amyloid nucleation *in vivo*^19–25^. Fluid flows in blood, cerebrospinal fluid, intramural periarterial drainage or interstitial systems are torrents at the microscopic scale, exerting various types of flow stresses including laminar-flow-dependent shear stress and extensional flow stress. The laminar-flow shear stress is governed by the shear rate (s^−1^), a gradient in velocity perpendicular to the direction of flow. Beyond a critical shear rate, a transition from diffusion-limited flow to advection dominated flow occurs, enabling shorter collision times between solutes and resulting in accelerated aggregation^25^. On the other hand, an extensional flow field is generated by a gradient in velocity in the direction of flow and is characterized by the strain rate (s^−1^). The extensional flow is enhanced when the channel diameter decreases or two channels merge. There are an increasing number of reports that extensional flows result in protein aggregation including amyloid fibrils^20,26,27^. Both laminar flow and extensional flow cause mechanical stresses in liquid, which are believed to trigger protein aggregation. However previous studies did not consider the role of supersaturation in fluid flow stresses. It is likely that the direct role of flow stresses is to break otherwise persistent supersaturation.

Here, during our challenge combining ultrasonication and microchannels, we found that peristaltic pump flow of hen egg white lysozyme (HEWL) resulted in amyloid formation. Peristaltic pump-dependent amyloid formation is common to varying disease-associated amyloidogenic proteins: α-synuclein (αSN), amyloid β 1-40 (Aβ40), and β2-microglobulin (β2m), suggesting that shear stress i*n vivo* triggers amyloid nucleation. We performed a numerical simulation to evaluate the stress in the liquid during the operation of the peristaltic pump and found that significantly large shear stress can be created in the liquid by the peristaltic motion of rotors in contact with the tubes, suggesting that shear stress is the key mechanical factor of amyloid nucleation *in vivo*. We designed a peristaltic pump-dependent amyloid inducer with fluorescence detection and imaging, which will be useful for assessing amyloidogenic risks^10,28^ and analyzing the kinetics of shear stress-dependent amyloid formation.

## Results

### Peristaltic pump-dependent amyloid formation of HEWL

We previously reported the ultrasonication- or stirring-dependent amyloid formation of HEWL at pH 2 and 2.0 M guanidine hydrochloride (GuHCl), where HEWL assumes a native fold although the stability decreased: the denaturation-midpoint was 3.0 M GuHCl^29–31^. We found that flowing of HEWL solution with the peristaltic pump under the same conditions at air-conditioned room temperature (20-25°C) resulted in a marked increase in amyloid-specific thioflavin T (ThT) fluorescence when the recovered solution was examined. The increase in ThT fluorescence suggested the formation of amyloid fibrils.

To monitor the ThT fluorescence increase in detail, we constructed a looped system (Extended Data Fig. 1), in which a peristaltic pump was used to flow the HEWL solution and a fluorometer with a flow cell was used to monitor ThT fluorescence. We used an ISMATEC peristaltic pump and a JASCO FP920 fluorometer. The ThT fluorescence increased when the applied HEWL came to the fluorometer for the first time (Fig. 1b). Then, the ThT peak appeared every 20 min, accompanied by increases in peak and baseline intensities. After several cycles, the ThT intensity saturated at a high level, suggesting a concomitant progression of amyloid formation and dilution within the loop. The ThT fluorescence monitored by the fluorometer showed a basal-level increase accompanied by sharp spikes. Excluding those evidently produced by air bubbles, spikes were likely to be caused by large ThT-positive aggregates.

**Fig. 1.**
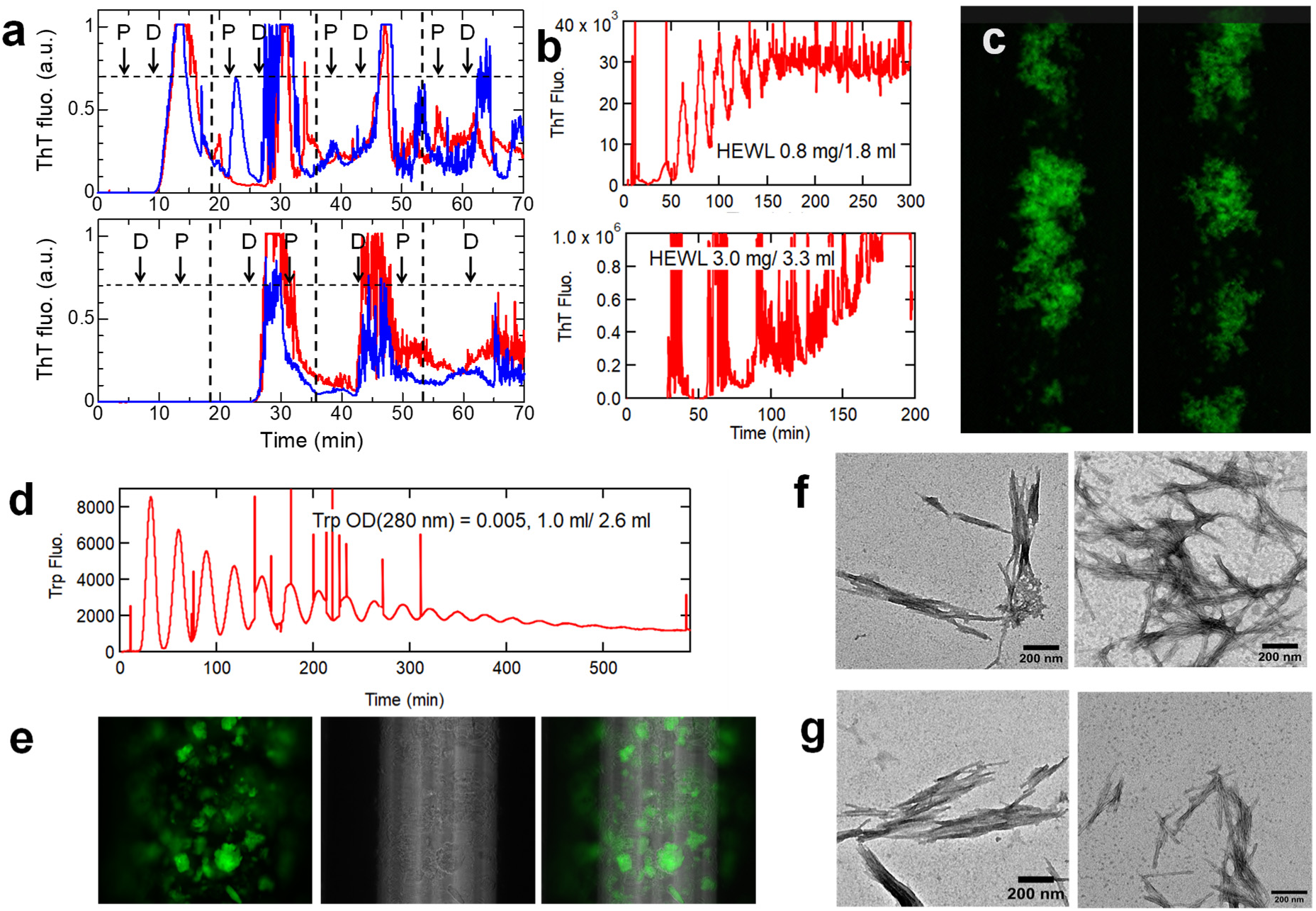
Peristaltic pump-dependent amyloid formation of HEWL. (a) The role of the peristaltic pump confirmed by exchanging the order of the peristaltic pump (P) and fluorescence detector (D). ThT fluorescence time-courses for alternative set-ups are shown. The HEWL amount applied was 0.375 mg, the total loop volume was 2.2 ml, and each loop cycle took 18 min. Red and blue curves show different experiments. (b) Representative ThT profiles at different HEWL amounts indicated. (c) ThT-fluorescence imaging of peristaltic pump-induced amyloid fibrils under flow. (d) The flow profile of tryptophan monitored at 350 nm. (e) Phase-contrast image of ThT-positive aggregates and its overlay on the ThT-fluorescence image. (f, g) TEM images of peristaltic pump-(f) or ultrasonication-induced amyloid fibrils (g).

In the present system, neither ultrasonication nor stirring was required. Among several possibilities by which HEWL formed ThT-positive aggregates (e.g., interactions with tube surfaces), the reaction trigger was attributed to the peristaltic pump. Because the peristaltic pump was located between the drawing inlet and fluorometer, the first passage of HEWL through the peristaltic pump was likely to trigger the reaction. To confirm the role of the peristaltic pump, we exchanged the positions of the peristaltic pump and fluorescence detector. With the exchanged set up, there was no ThT fluorescence increase during the first passage of HEWL to the detector, demonstrating the role of the peristaltic pump (Fig. 1a). Subsequent experiments were performed with the peristaltic pump located before the fluorescence detector.

As a control, we flowed a tryptophan (i.e., N-acetyl L-tryptophan-amide) solution and monitored its fluorescence at 350 nm (Fig. 1d). The tryptophan fluorescence showed repeated peaks with decreasing maximal and increasing minimal intensities, showing flow-dependent monotonous dilution within the loop.

We then installed a fluorescence microscope to simultaneously monitor the morphology of ThT-positive products (Fig. 1c, Extended Data Fig. 1, Sup. Video 1). Various sizes of ThT-positive aggregates up to tens of micrometers with clustered fluff-like morphology flowed one after another, followed by a period of darkness, consistent with repeated peaks monitored by ThT fluorescence. We observed the flow-dependent tumbling and change in morphology: venting and extension of fibrous clumps. These images revealed a shear flow with varying shear rates: faster at the center than at the near-wall position of the tube. The peristaltic flow stopped for 0.1 s every 4.6 s at a flow rate of 0.1 ml/min, leading to the flow-and-stop motion of aggregates. Once the reaction monitored by ThT fluorescence ended, aggregate morphology was considered mature; it was retained during subsequent cycles of flow.

We also measured phase-contrast bright mode images of these aggregates, showing that the regions with strong ThT-fluorescence overlapped with entire aggregates detected in the phase-contrast mode (Fig. 1e). This indicated that most of the aggregates formed under these conditions were ThT-fluorescence-positive.

To confirm that ThT-positive aggregates are amyloid fibrils, the solutions were recovered from the loop, and transmission electron microscopy (TEM) (Fig. 1f) and circular dichroism (CD) measurements (Extended Data Fig. 2a) were performed. TEM revealed a typical needle-like amyloid morphology with a diameter of approximately 10 nm and length up to several μm. The CD spectra showed a β-sheet-dominated structure but with a small intensity. We later found that, because of notable absorption to tube surfaces, the CD spectra of recovered solutions do not represent the secondary structures of reaction products.

Because the extensional flow apparatus with two syringes connected by a single capillary was reported to induce amyloid fibrils of β2m^20^, we flowed the HEWL solution with a syringe-pump repeatedly (Extended Data Fig. 3), where the flow rate was set to be comparable with the standard flow rate (0.1 ml/min) used for peristaltic pump experiments. After passage through the fluorescence detector, the recovered eluent (2.2 ml) was used to fill the syringe with a 4-way cock and flowed again. Although the repeated rounds of syringe-pumping slightly increased the ThT fluorescence, it was at most 5% of the peristaltic pump-dependent ThT burst, indicating that the peristaltic pump is highly effective in inducing amyloid fibrils.

However, during the peristaltic pump experiments, we noticed that the recovered solutions did not show the yellow opal color typical of ultrasonication-induced amyloid fibrils. We compared the peristaltic pump-induced amyloid fibrils with those created by ultrasonication^18,29,30^. When the monomer HEWL solution in a cell of the fluorometer was irradiated with the attached ultrasonicator^18^, ThT fluorescence increased cooperatively, consistent with previous reports^29,30^ (Extended Data Fig. 4a). Under quiescent conditions, no increase in ThT fluorescence occurred within 24 h. Stirring of the solution by a stirrer increased the ThT fluorescence intensity slightly.

Even considering the dilution that occurred during peristaltic pump experiments, the specific ThT fluorescence intensities at a constant protein concentration were much lower (approximately 10%) than those of the ultrasonication-induced HEWL amyloid fibrils (Supporting Table 1). It was likely that the absorption of amyloid fibrils to loop inner surfaces including the flow cell reduced the recovery yield, making the quantification of amyloid fibrils difficult. In fact, when the preformed HEWL amyloid fibrils were flowed, we observed a marked decrease in ThT fluorescence during flow cycles (Extended Fig. 5), indicating the absorption of preformed amyloid fibrils to the loop surfaces.

We then examined the dependence of the ThT fluorescence kinetics on the amount of HEWM monomers applied (Extended Fig. 6a, and c, Supporting Table 2), showing the complicated dependence, which is likely to be caused by a combined effect of the intense triggering of amyloid nucleation and absorption of amyloid fibrils to loop surfaces.

### Different types of peristaltic pumps

We used different types of peristaltic pumps (ATTO and AS-One peristaltic pumps) to examine whether amyloid formation is common to peristaltic pumps. When examined with the same HEWL solution, other peristaltic pumps also showed amyloid formation (Fig. 2). On the other hand, the ThT-fluorescence kinetics differed depending on pumps. ISMATEC and ATTO peristaltic pumps have 12 and 6 rotors, respectively, and AS-One pump has one rotor whose position rotates. The flow cycles visualized by the preformed HEWL amyloid fibrils were distinct (Sup. Video 2): The ISMATEC pump showed cycles of 4.5-s flow and 0.1-s stop, ATTO pump showed the cycles of 26-s flow and 4-s stop, and AS-One pump showed the cycles of 27-s flow with gradual slow down and 3-s stop. In addition, the tube sizes in the pump were different, although we used the same tube out of the pump (inner diameter: 1.0 mm; length: ∼2.0 m). Inside peristaltic pumps, ATTO and As-One pumps have wider tubes (inner diameter: 2.0 mm) and the ISMATEC pump used the tube with the same inner diameter (1.0 mm) as the main loop. These differences were responsible for the variation in the ThT fluorescence pattern.

**Fig. 2.**
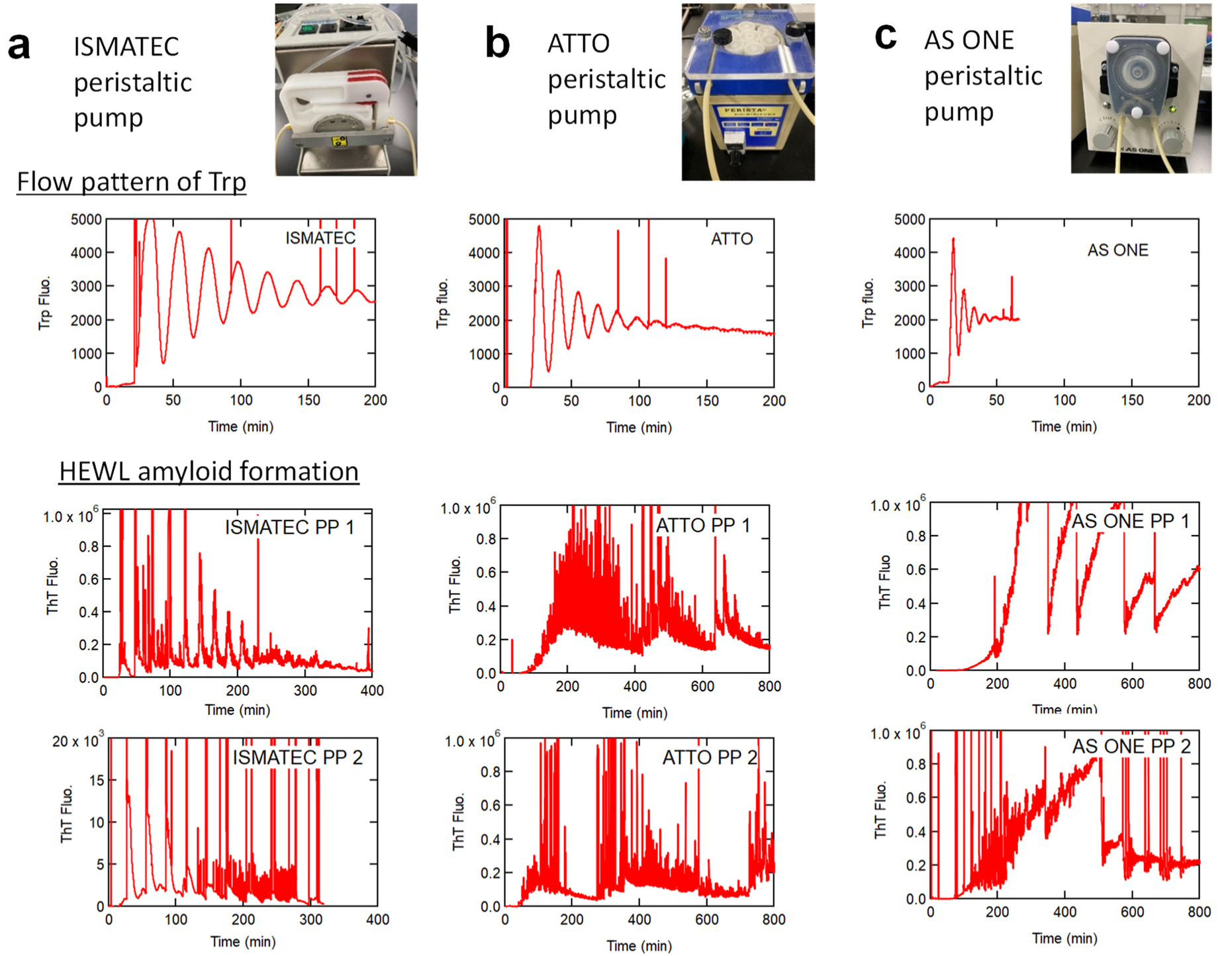
Comparison of (a) ISMATEC, (b) ATTO, and (c) AS-One pelistaltic pumps. Photos, flow patterns with N-acetyl-tryptophan-amide monitored at 350 nm, and examples of the kinetics for HEWL amyloid formation. For the ISMATEC peristaltic pump, the HEWL solution at 1.0 mg/ml was applied for 10 min at 0.1 ml/min. Similar HEWL amounts and conditions were used for ATTO and AS One pumps.

Intriguingly, the AS-One peristaltic pump exhibited a cycle of a gradual increase and sudden decrease in ThT fluorescence (Fig. 2b). It was likely that the gradual slow down and 3-s stop allowed deposition of amyloid fibrils on the surface of the flow cell, which was peeled off after a certain period of accumulation time.

### α-Synuclein

To examine the generality of peristaltic pump-dependent amyloid formation, we first used αSN, an intrinsically disordered protein of 140 amino acid residues associated with synucleinopathies, including Parkinson’s disease, dementia with Lewy bodies, and multiple-system atrophy^6^. αSN forms amyloid fibrils at high concentrations of salts at a neutral pH^32^. Ultrasonication-dependent amyloid formation of αSN occurred at pH 7.0 in the presence of 0.5 M Na_2_SO_4_ (Extended Data Fig. 4b). With the peristaltic pump system, we typically applied 1.0 mg of αSN (0.5 mg/ml αSN at a flow rate of 0.1 ml/min for 20 min). With the fluorometer, a marked increase in ThT fluorescence was observed upon the first passage of the applied solution (Fig. 3a). As was the case with HEWL, fluorescence microscopy revealed clustering fluff-like aggregates with shear flow-dependent dynamic motions, including tumbling and deformation (Fig. 3b, Sup. Video 1).

**Fig. 3.**
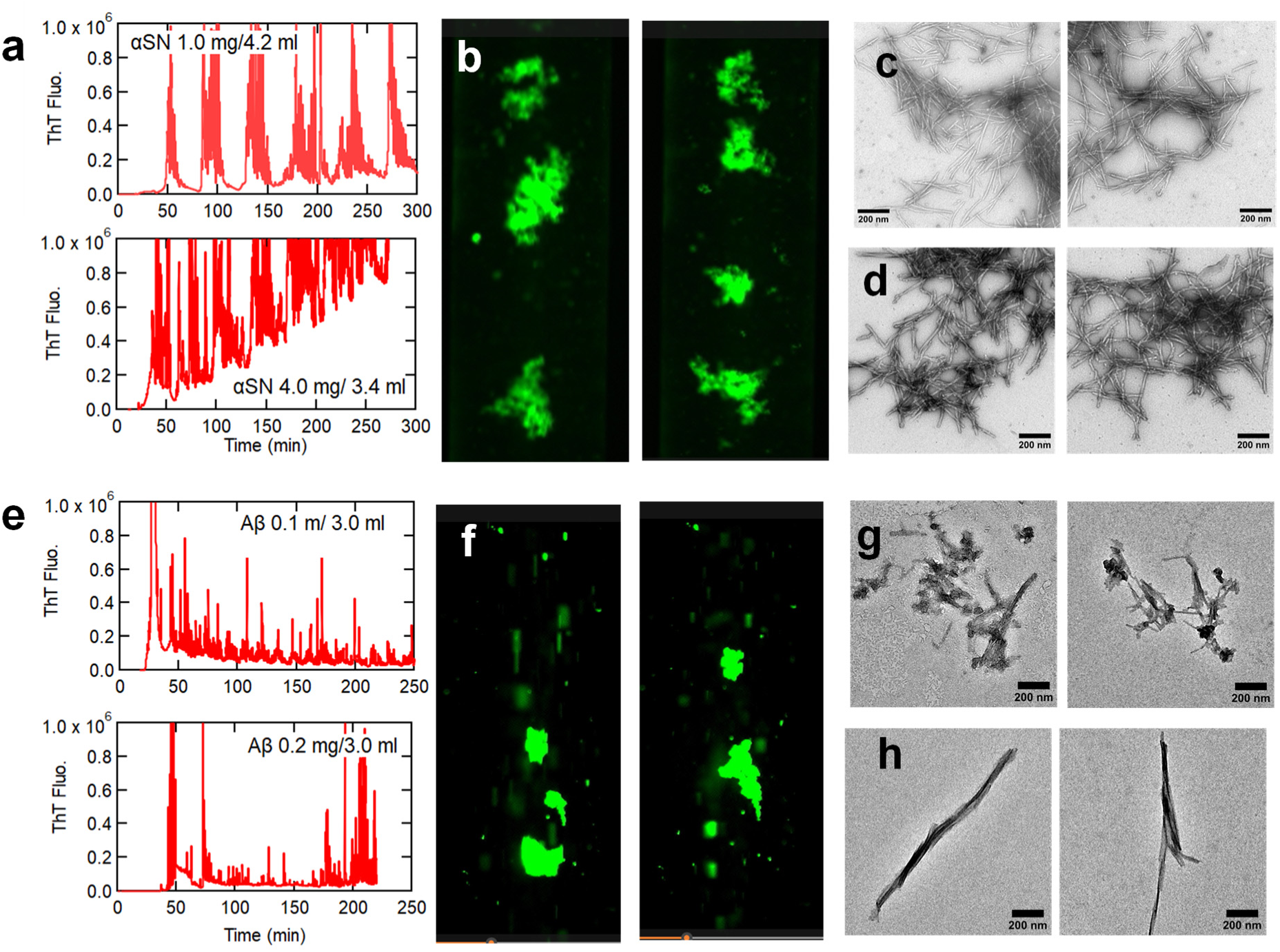
Peristaltic pump-dependent amyloid formation of αSN (a-d) and Aβ40 (e-h). Typical profiles of amyloid formation observed at 485 nm (a, e), fluorescence microscopy images (b, f), and TEM images (c, g) are shown. TEM images of amyloid fibrils prepared by ultrasonication (d, h) are also shown.

TEM of recovered solutions showed clear amyloid fibril images (10 nm in width and several μm in length) (Fig. 3c). In contrast, the CD spectra often did not exhibit a typical β spectrum (Extended Data Fig. 2b). A comparison of ThT fluorescence values for those produced by ultrasonication and the peristaltic pump indicated that the apparent amount of amyloid formed by the latter was less than that by the former (Supporting Table 1) because of the absorption of amyloid fibrils.

We studied the dependence of ThT fluorescence kinetics on the amount of αSN applied (Extended Data Fig. 6b, d). The results were similar to those of HEWL, indicating that the peristaltic pump triggered amyloid nucleation in the first round even when the total amount applied was low and subsequent kinetics were complicated by a combination of amyloid growth and absorption.

### Amyloid β40 peptide

The peristaltic pump also triggered amyloid formation of Aβ40, associated with Alzheimer’s disease, upon the first passage when monitored by ThT fluorescence (Fig. 3e). Subsequent cycles decreased the peak intensity because of absorption to tube surfaces, which was more serious than the cases of HEWL or αSN. Fluorescence microscopy showed the flowing of large amyloid clumps under shear flow (Fig. 3f, Sup. Video 1, 3). Self-association of preformed amyloid clumps and their absorption to the flow-cell surface with subsequent peeling off were observed more often than with other proteins, indicating the sticky nature of Aβ40 fibrils. Amyloid clumps were often too large to tumble and were about to occlude the channels; this happened for some experiments and are likely to occur *in vivo*. Upon ‘unintentional’ flicking of the tube after the apparent calming down of amyloid ‘river’, many amyloid clumps suddenly surged, showing the detaching of absorbed amyloids (Sup. Video 3). Subsequent ‘intentional’ flicking indeed caused the expected surge in amyloid fluffs. These debris flow, if happened *in vivo*, will be irreversible, producing serious damages including occlusion of *in vivo* channels. We name the video ‘Amyloid River of No Return’ after the 1954 motion picture ‘The River of No Return’.

Cerebral amyloid angiopathy is characterized by cerebrovascular Aβ deposition with a high Aβ40/42 ratio: mainly Aβ40 forms amyloid deposits on vascular wall leading to acute disruption in the intramural perivascular drainage (or intramural periarterial drainage) flow and cerebral blood flow^33–35^. The fluorescence microscopy videos imply early stages of cerebral amyloid angiopathy. It is noted that, in our experiments, Aβ40 deposited on the inner surface of tube, while, in cerebral amyloid angiopathy, Aβ40 deposits on the external surface of vascular walls.

TEM showed rigid amyloid morphology with a diameter of 10 nm and length of 1-2 μm (Fig. 3g). CD spectra showed conversion from disordered monomers to a β-sheet structure typical of amyloid fibrils (Extended Data Fig. 2c), although the precise evaluation was difficult because of amyloid absorption to loop surfaces.

We also examined the ultrasonication-induced amyloid formation of Aβ40 monitored by a fluorescence spectrophotometer (Extended Data Fig. 4c). The ultrasonication induced amyloid fibrils effectively and the specific ThT fluorescence value was higher than that induced by the peristaltic pump (Supporting Table 1). All these results showed the peristaltic pump-triggered efficient formation of Aβ40 amyloid fibrils and their marked propensity for self-association and absorption.

### β2m

Finally, we used β2m, responsible for dialysis-related amyloidosis^17,36^. β2m has been reported to form amyloid fibrils at pH 2 with a moderate concentration of salt (e.g., 0.2 M NaCl) under stirring or ultrasonication^31,37,38^. Although amyloid formation at a neutral pH, where patients develop amyloid deposits, was previously difficult, several conditions have been explored to induce amyloid fibrils including the presence of low concentrations of SDS^39^ or polyphosphate^40^, or at a high temperature combined with solution agitation^17,38^. However, amyloid formation at a physiological pH and temperature without additives remains challenging.

We first examined the β2m amyloid formation at pH 1.8 in 0.4 M NaCl (Fig. 4a). ThT fluorescence increased upon the first passage through the peristaltic pump and fluorescence microscopy revealed fluff-like aggregates flowing and tumbling with shear-dependent dynamic motions (Fig. 4d, Sup. Video 1). Amyloid formation was confirmed by TEM (Fig. 4h). CD spectroscopy showed the spectrum of the β-sheet although the intensity was reduced because of absorption to tube surfaces (Extended Data Fig. 2d). As for other proteins, specific ThT intensity of recovered solution after the peristaltic pump treatment was low because of absorption (Supporting Table 1).

**Fig. 4.**
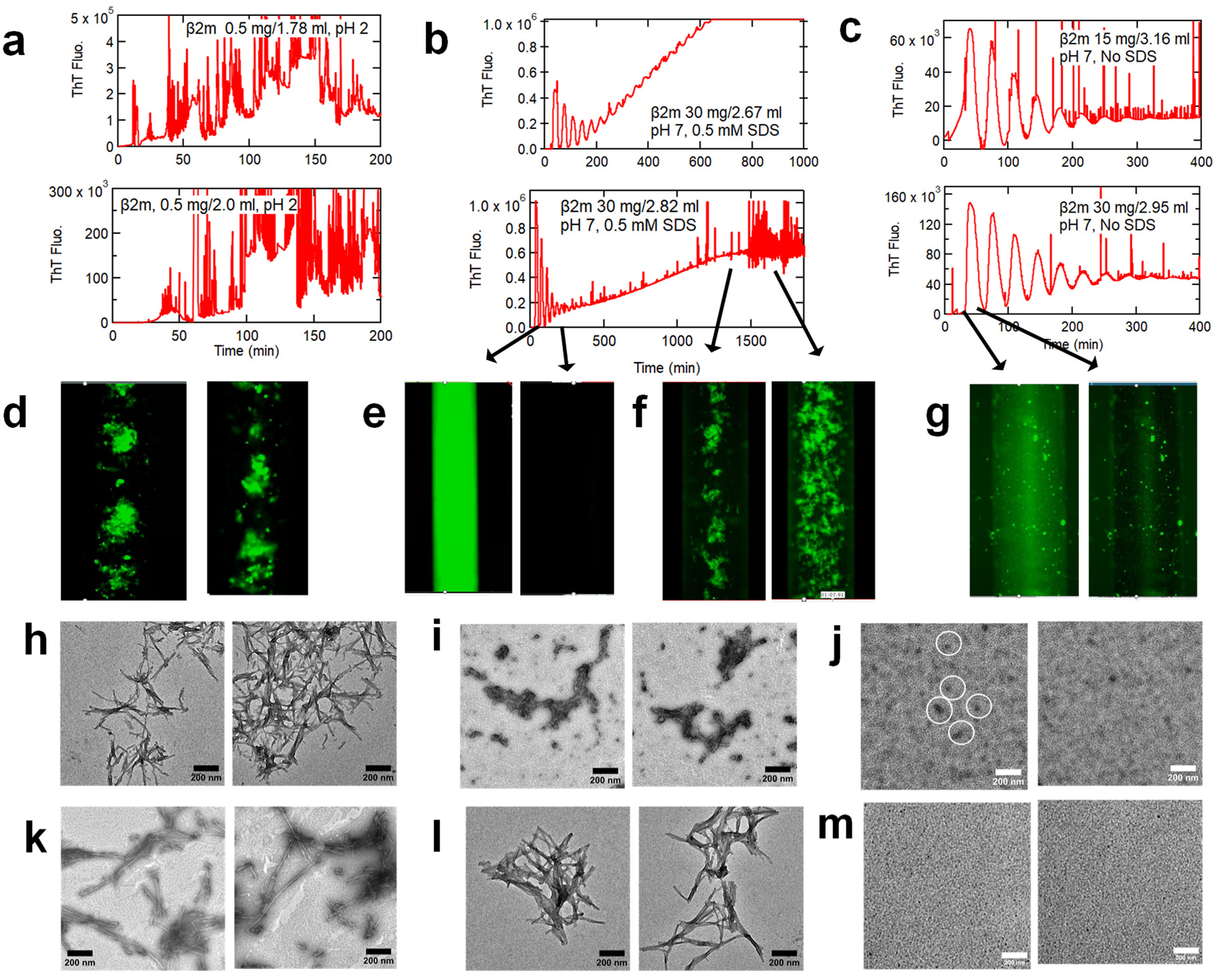
Peristaltic pump-dependent amyloid foramtion of β2m at 20 mM HCl and 0.4 M NaCl (a, d, h, k), at pH 7.0 in the presence (b, e, f, i, l) or absence (c, g, j, m) of 0.5 mM SDS. Typical profiles of amyloid formation observed at 485 nm (a-c), fluorescence microscopy images (d-g), and TEM images (h-j) are shown. In the left panel of (j), oligomers are indicated by circles. TEM images of amyloid fibrils prepared by ultrasonication are also shown (k-m).

At pH 7.0 in the presence of 0.5 mM SDS, the peristaltic pump revealed intriguing two-step amyloid formation monitored by ThT fluorescence (Fig. 4b). The first passage of β2m through the peristaltic pump markedly increased the ThT fluorescence. Subsequent cycles resulted in oscillating peaks with a decreasing intensity because of dilution. The fluorescence microscopy movie at a high playback speed (x256) looks like a firefly flashing (Fig. 4e, Sup. Video 4). We observed no obvious aggregate or fluff at the fluorescence microscopy resolution. Several hours later, ThT fluorescence started to increase again but gradually, producing fluff-like aggregates. The fluff-like aggregates were confirmed to be amyloid fibrils by TEM (Fig. 4i). As for other proteins, CD or ThT fluorescence of recovered solutions was not accurately evaluated because of absorption to tube surfaces (Extended Data Fig. 2e, Supporting Table 1). However, the ThT fluorescence value after the end of flashing (0.15 x 10^8^) was approximately 25% of that of amyloid fibrils (0.6 x 10^8^), suggesting that the intermediates are partly amyloid-like (Fig. 4b, Sup. Video 4). Although the recovered solution of the intermediates did not show the notable ThT intensity, it was likely that the absorption decreased the recovery (Sup. Table 1).

When the peristaltic pump effects were examined in the absence of SDS at pH 7.0, only the first step (firefly flashing) occurred (Fig. 4c, g, Sup. Video 5), suggesting that, even in the absence of SDS, the intermediate state with notable ThT fluorescence accumulated. In the absence of SDS, nothing happened even under ultrasonication (Extended Data Figs. 4, 7). TEM images of the recovered β2m solution showed no fibrillar images. Instead, small aggregates, resembling so-far reported oligomers^41–45^, were observed (Fig. 4j). Importantly, we previously analyzed the SDS-dependent first-step product by analytical centrifugation, reporting that they are oligomers consisting of approximately five β2m monomers^46^. The SDS-induced oligomers showed a CD spectrum of partly disordered conformation, consistent with the present results (Extended Data Fig. 2f). The SDS-induced β2m oligomers were useful to promote seed-dependent amyloid growth under neutral pH conditions^39^.

We separated the effects of SDS and ultrasonication with the fluorescence spectrophotometer (Extended Data Fig. 7). In the presence of 0.5 mM SDS at pH 7.0, the SDS-dependent first step occurred under stirring with a time constant of approximately 30 min at 0.5 mg/ml β2m and depended on the β2m concentration: faster and slower at higher and lower β2m concentrations, respectively. The second step leading to amyloid fibrils required ultrasonication. In contrast to SDS-dependent oligomer formation, peristaltic pump-dependent oligomer formation was independent of the β2m concentration, indicating the strong power of the peristaltic pump.

Here, the ThT-positive oligomers detected by the peristaltic pump may be an off-pathway trapped intermediate of amyloid formation (Extended Data Fig. 7a). Although the differentiation of on- and off-pathway intermediates is usually not straightforward^47^, the ultrasonication-dependent full conversion of oligomers to amyloid fibrils supported that they are off-pathway trapped intermediates. The accumulation of off-pathway intermediates can be explained by Ostwald’s ripening rule of crystallization, according to which the morphologies of crystals change over time, guided by their kinetic accessibilities and thermodynamic stabilities^48^. Following this rule, the ThT-positive oligomers are rapidly formed as trapped intermediates, which can be converted to more stable and structured amyloid fibrils. Taken together, even in the absence of SDS at pH 7.0, prolonged exposure of β2m to shearing stresses increases the risk of amyloid formation.

## Discussion

### Finite-element-method (FEM) simulation

To evaluate the mechanical stimuli induced in the flowing solution by the peristaltic pump, we performed two-dimensional finite-element-method (FEM) simulation with COMSOL Multiphysics ver. 6.2 by combining the solid mechanics and fluid mechanics calculations (Fig. 5). The model consists of two rubber sheets with a thickness of 1 mm and length of 175 mm, with a 1-mm thick water layer between them (corresponding to the rubber tube with outer and inner diameters of 3 and 1 mm, respectively), and 3-mm steel rotors located above the rubber sheets. The rotors were first pressed vertically against the rubber sheet to decrease the solution-layer thickness, horizontally moved along the flow direction, and then stopped. During the rotor movement, the shear rate and then shear stress in the solution were calculated.

**Fig. 5.**
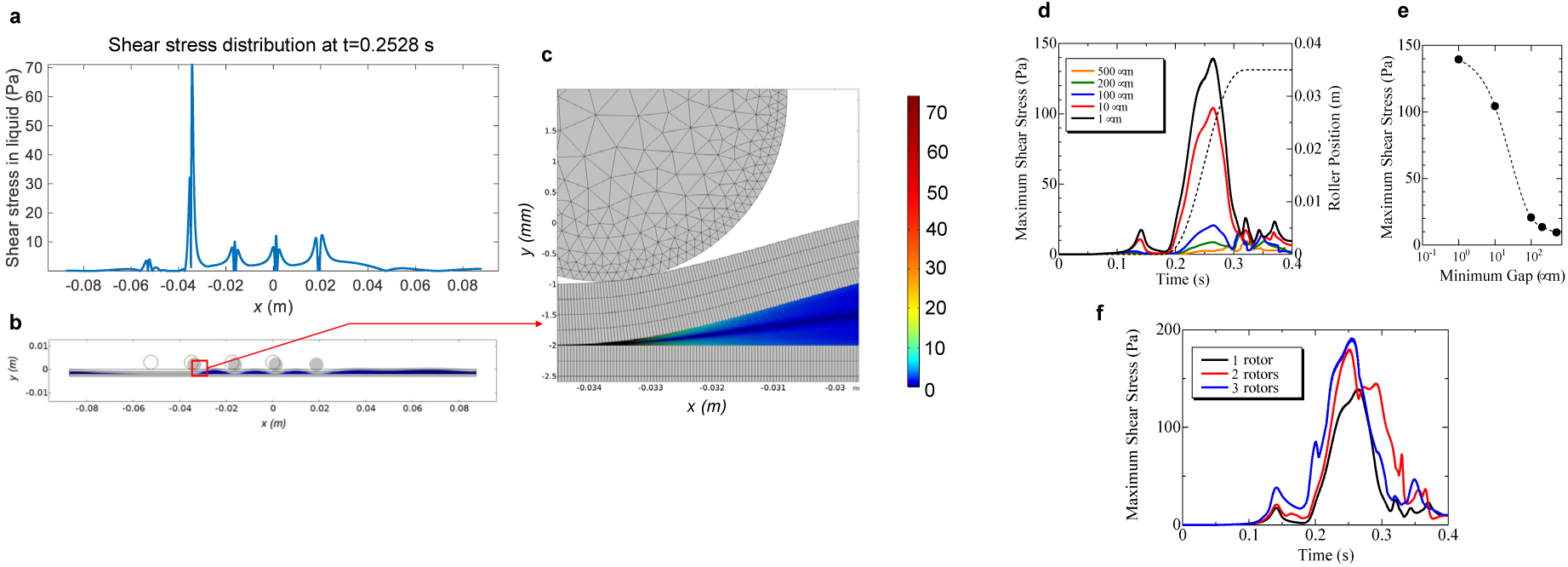
Numerical analysis of shear stress induced in liquid by movement of a contacting rotor. The behavior of the tube filled with solution was simulated by a two-dimensional model where the two rubber sheets accomodate water in the gap between them. The thickness and length of the rubber sheets were 1 and 175 mm, respectively, and the initial gap was 1 mm. The 3-mm-diameter steel rotor was first pressed vertically against the rubber sheet, moved horizontally along the flow direction, and stopped, while the shear rate and shear stress were calculated by FEM analysis. The maximum and minimum mesh sizes were 2.5 and 0.2 μm, respectively, and the time step was 0.1 ms. The rubber sheets were fixed at both ends, and the open boundary condition was adopted for liquid. To allow for large deformations, moving mesh was used. (a) A snap shot of shear-stress distribution during the movements of four rotors, (b) corresponding rotor positions, and (c) an enlarged view near the rotor located most downstream. These are results for the case with a remaining gap of 50 μm. (d) Change in the maximum shear stress with single-rotor movement in an axial direction for various minimum gaps at the first roller pushing action. The broken line denotes the rotor position, indicating that the shear stress is maximal near the maximum rotor speed. (e) Dependence of the maximum shear stress on the minimum gap for the single-rotor case. (f) Dependence of the maximum shear stress on the number of rotors.

Supplementary Video 6 shows the change in the shear-stress distribution in the liquid layer during the rotor movement when 1, 2, and 4 rotors was used. Significantly large shear stresses were locally produced, which must be the cause of the peristaltic pump-dependent amyloid nucleation. The laminar flow-dependent maximum shear stress in flowing liquid can be evaluated by 4μU_ave_/R for a flow channel with circular cross-section, where μ, U_ave_, and R denote the solution viscosity, average flow velocity, and inner radius of the flow channel, respectively. The average flow velocity in our experiments was typically 0.002 m/s, yielding a maximum shear stress of 0.02 Pa or less for the 1-mm inner diameter tube. Therefore, the rotor movement can cause a much larger shear stress than the laminar flow.

Figure 5 indicates that the shear stress in solution can be increased by decreasing the thickness of the remaining solution layer in the first rotor push action. Because the FEM calculation failed when the remaining solution layer was too thin, we made the calculation with a remaining layer thickness greater than 1 μm. In the actual peristaltic-pump operation, the solution layer can be nearly zero and shear stress is estimated to be larger. In addition, the shear stress can be further increased by increasing the number of rotors (Fig. 5c), reaching nearly 200 Pa, a 10,000-fold increase compared with simple laminar flow.

To induce a large shear stress with laminar flow, we need to use a flow channel with very small cross-section. For example, a shear stress of ∼100 Pa can be locally achieved using a microchannel with a cross-section of 80 μm x 62 μm^49^. However, aggregates containing rigid amyloid fibrils will quickly occlude such a narrow channel, making the flow assay impractical. Therefore, the moving motion of the rotor in contact with the tube in the peristaltic pump is a rare phenomenon that can generate very large shear stresses, even in channels with large inner diameters.

### Kinetic analysis

We propose a theoretical model for understanding the amyloid-formation behavior in the flow assay composed of the peristaltic pump and fluorescence detector (Methods), in which the concentrations of the applied monomer (*C_M_*) and product (fibrils) (*C_F_*) vary not only with the progression of the chemical reaction (amyloid formation), but also with their self-diffusion and with the axial convection due to the flow. Concerning the chemical reaction, we consider the two-step fibril-formation model, which is characterized by the reaction-velocity constant for the primary nucleation, *k_n_*, and that for the fibril elongation, *k_g_*^50,51^.

Figure 6 compares simulation calculations with some experiments. Although there are some ambiguous parameters, such as diffusion coefficients of monomers and fibrils, the analytical model reproduces the experimental results and is able to explain fibril-formation under convection flow. However, it failed to simulate some experiments when the initial monomer amount was small, as shown in Fig. 6c. In the experimental results, certain amounts of fibrils were produced from the beginning because of the strong amyloid-inducing capacity, but in the simulation, the amount of fibrils increased gradually. In the experiments, absorption of preformed fibrils reduced the flowing fibrils. Moreover, this discrepancy may be attributed to its spatiotemporal localization. Even if fibrils are generated, their localization prevents further immediate propagation, as occurring in the test tube.

**Fig. 6.**
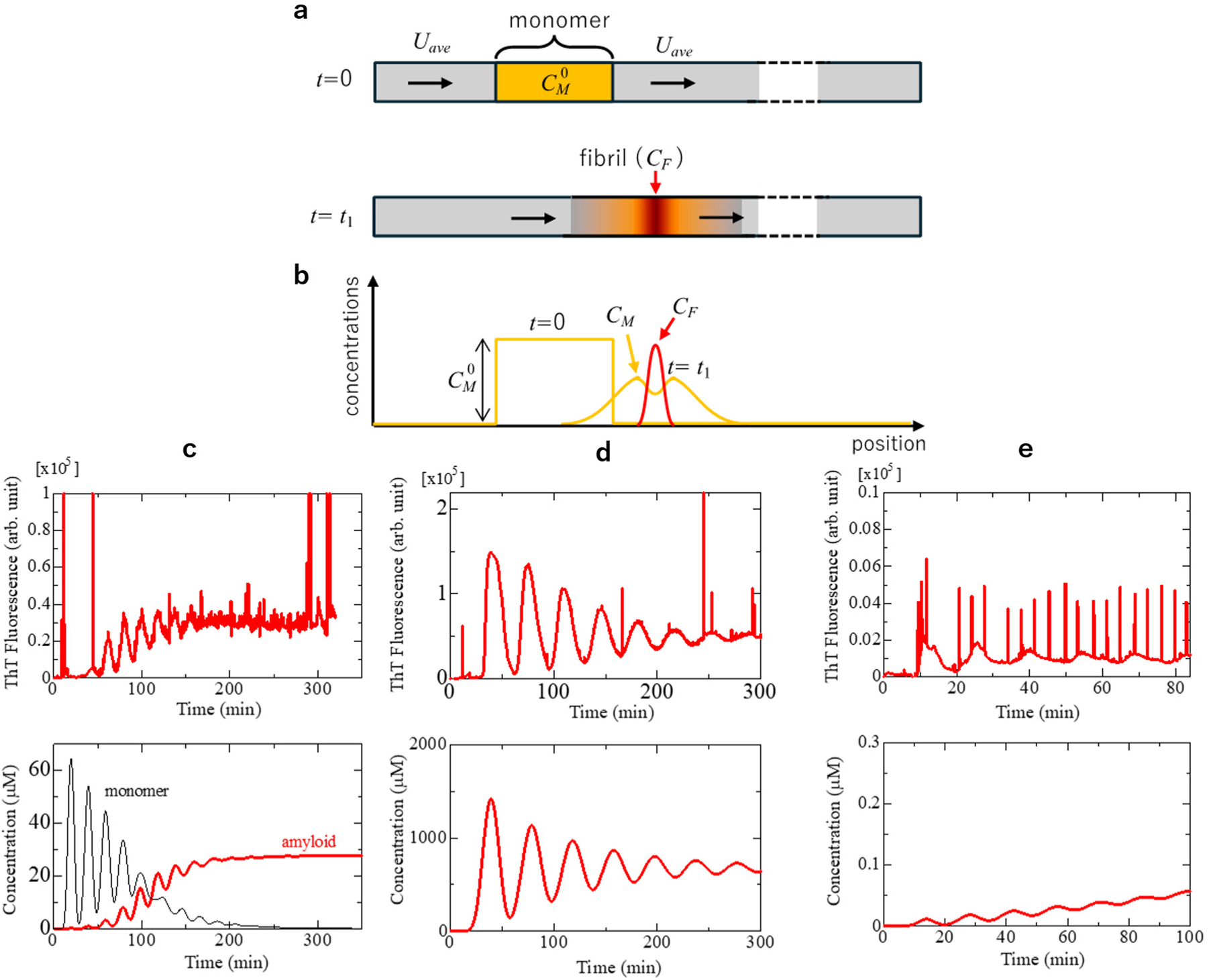
Simulation of peristaltic pump-dependent amyloid formation. (a) One-dimensional model for calculating monomer and fibril concentrations under axial flow. (b) Schematic of time variation of the concentration distributions. (c-e) Comparison between experiments (upper) and numerical simulations (lower) for (c) 1.0 mg/mL HEWL, (d) 20 mg/mL β2m, and (e) 0.5 mg/mL HEWL. The protein concentrations are those of applied solution and the total amounts applied were (c) 0.8 mg, (d) 30 mg, and (e) 0.25 mg. The parameters used are as follows. Legth of tube: 2.5 m; diffusion coefficinets of monomer (*D_M_*) and fibril (*D_F_*): 7×10^-10^ and 2×10^-10^ m^2^/s, respectively; reaction-velocity constants for nucleation (*k_n_*) and fibril growth (*k_g_*): 1×10^-6^ s^-1^ and 3×10^-5^ s^-1^μM^-1^, respectively; flow velocity: (c) 100, (d) 50, and (e) 140 μl/min.

### Perspectives

Supersaturation is a prerequisite to form amyloid fibrils as well as crystals. According to solubility and supersaturation-limited amyloid formation^10–15^, in the metastable region of supersaturation located slightly above the solubility limit, supersaturation persists in the absence of seeding. Thus, triggers based on mechanical stimuli are essential for breaking supersaturation. Although fluid-flow and associated shear stresses have been increasingly focused on^20–25^, supersaturation has not been addressed in the context of shear stresses. We propose that the direct contribution of fluid-flow stresses is the mechanical breaking of supersaturation. Once supersaturation is broken, the thermodynamic equilibrium of soluble monomers and amyloid fibrils becomes established^10,17,31^.

We showed that peristaltic pumps are unexpectedly powerful for triggering amyloid formation of proteins in general. In the current loop system, the absorption of preformed amyloid fibrils to loop inner surfaces including the flow cell complicates the observed kinetics. However, this complexity mimics amyloid deposition *in vivo,* where various tissues and organs are connected by fluid flows including blood, cerebrospinal fluid, interstitial fluid and intramural perivascular drainage systems. Although no further supply of amyloid proteins occurs in our loop system, the continuing supply of monomeric proteins will lead to the massive growth of amyloid deposits in specific tissues or organs near the triggering sites. Markedly sticky Aβ images may mimic early stages of cerebral amyloid angiopathy (Sup. Video 3)^33,34^. The most serious concern from our ‘Amyloid River of No Return’ is the occulusion of *in vivo* channels, which will be hard to be returned. The firefly flashing β2m oligomers might occur in patients under dialysis therapy where peristaltic pump-based hemodialysis machines are used.

Finally, the market effectiveness in spite of a concise setup will make the peristaltic pump system useful for early-stage diagnosis by monitoring susceptibility risk biomarkers even without seeds^28^. From the viewpoint of supersaturation-limited amyloid formation, therapeutic strategies for reducing the degree of supersaturation (Equation (1)), a measure of the risk of ‘amyloid ignition’, will be more effective than extinguishing the ‘amyloid fire’. Such an approach has been undertaken by the ultrasonication-dependent amyloid inducer, a HANABI system, seeking a high throughput analysis of the risk of amyloid ignition with a microplate reader^10,17^. The current peristaltic pump system might bring an alternative approach, enabling the imaging and kinetic analysis of the shear stress-forced amyloid ignition in real time and at a microliter scale. These will contribute to realizing the return of amyloid river like the ending of “River of No Return”.

## Methods

### Proteins and chemicals

Recombinant human αSN was expressed in *Escherichia coli* and purified as described previously^52^. Recombinant human β2m with an additional methionine residue at the N terminus was expressed using *Escherichia coli* and purified as described previously^53^. HEWL^30^ and Aβ40^32^ were purchased from Nacalai Tesque (Kyoto, Japan) and Peptide Institute Inc. (Osaka, Japan), respectively. The fluorescence dye ThT was obtained from Wako Pure Chemical Industries (Osaka, Japan). N-Acetyl L-tryptophan amide and other reagents were purchased from Nacalai Tesque.

### Peristaltic pump-dependent amyloid formation

HEWL, αSyn and β2m were dissolved in deionized water. Aβ40 was dissolved in 0.2% (w/w) ammonia solution. Then, the following buffers were used to enable the solution to flow by the peristaltic pump: HEWL in 20 mM HCl (pH 1.8) and 2.0 M GuHCl; αSN in 50 mM sodium phosphate buffer (pH 7.0) and 0.5 M Na_2_SO_4_; Aβ40 in 20 mM sodium phosphate buffer (pH 7.0) and 0.05 M NaCl; β2m under acidic conditions in 20 mM HCl (pH 1.8) and 0.4 M NaCl; β2m under neutral pH conditions in 50 mM sodium phosphate buffer (pH 7.0) and 0.1 M NaCl in the presence or absence of 0.5 mM SDS. All protein solutions contained 5 μM ThT. Experiments were performed at an air-conditioned room temperature (i.e. 20-25 °C).

The assays with peristaltic pumps were carried out with the looped system (Extended Data Fig. 1). We used an ISMATEC peristaltic pump (REGLO Analog MS-2/12) and a JASCO FP920 fluorometer with a16-μl flow cell volume. These apparatuses were connected by a silicone tube with an inner diameter of 1.0 mm. Typically, 0.8 ml of a 1.0-mg/ml HEWL solution (0.8 mg HEWL) was drawn at a flow rate of 0.1 ml/min and then the inlet and outlet were connected to form a loop. The total length of the loop was approximately 1.5 m with a total volume of 2-3 ml. To measure the precise volume, we recovered the solution after the experiments and weighed it. The recovered solutions were also used for CD, TEM, and ThT fluorescence measurements.

We also used different types of peristaltic pumps (i.e. Model AC-2120 (ATTO Co., Tokyo, Japan) and Model TP-10SA (AS-One Co., Osaka Japan) peristaltic pumps) to examine the generality of peristaltic pump-dependent amyloid formation (Fig. 2, Sup. Video 2).

An intelligent fluorescence detector (Model FP-920, Jasco Co., Ltd., Tokyo, Japan) with a PC integrator ChromNAV Lite was used to monitor the ThT or Trp fluorescence in the looped system. The flow cell volume was 16 μl. The excitation and emission wavelength were 445 and 485 nm, respectively, for ThT fluorescence and were 280 and 350 nm, respectively, for Trp fluorescence.

A fluorescence microscope (Model BZ X810, Keyence Co., Ltd., Osaka, Japan) was installed to simultaneously monitor the morphology of ThT-positive products (Fig. 1, Extended Data Fig. 1, Sup Video 1). Flow in a glass tube with a diameter of 1.2 mm was visualized at a low magnification (x 2). We used the BX filter GFP (excitation at 470 nm and emission at 495 nm) to monitor ThT fluorescence. We also measured the phase contrast images of aggregates to check the overlap with ThT fluorescence images.

We examined the dependence of amyloid formation on the amount of HEWL or αSN applied (Extended Fig. 6). For the current system employing the sample drawing, changing the apply volume of the same stock solution was more convenient than preparing solutions at different protein concentrations.

### Ultrasonication-dependent amyloid formation

Ultrasonication-dependent amyloid formation was measured with the Hitachi fluorescence spectrophotometer F7000 (Tokyo, Japan), in which the sample solution in a glass cuvette was irradiated with ultrasonic pulses from an ultrasonic generator tightly attached to the sidewall of the cuvette (Elekon Science Co., Chiba, Japan), as reported previously^18,54^.

### CD and TEM measurements

Far-UV CD spectra (approximately 200–250 nm) were obtained by the model J-820 spectropolarimeter (Jasco Co., Ltd., Tokyo, Japan) at 20 °C using a quartz cell with a 1-mm path length. CD data are expressed as the mean residue ellipticity. To calculate the mean residue ellipticity, the initial protein concentration of the sample solution was used. This obviously made it difficult to convert to molar ellipticity precisely because a significant portion of the protein might be absorbed on the inner surfaces of flow loops.

TEM images were obtained with a transmission electron microscope (JEOL JEM-1400). A 3-μL aliquot of the sample solution was placed on a collodion-coated copper grid (Nisshin EM Co.) for 30 s, and the remaining solution was removed with filter paper. The sample was then stained with a 1% (w/v) uranyl acetate solution for 30 s. Finally, the surface of the grid was rinsed with deionized water. TEM observation was performed with an acceleration voltage of 80 kV.

### Theoretical analysis of amyloid formation in the looped flow system

We considered the two-step fibril-formation model, which is characterized by the reaction-velocity constant for primary nucleation, *k_n_*, and that for the fibril elongation, *k_g_*^50,51^. The governing equations are then given by:

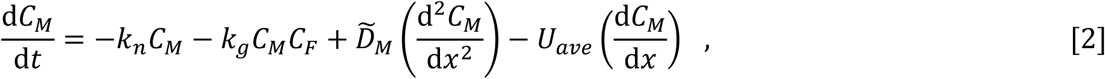

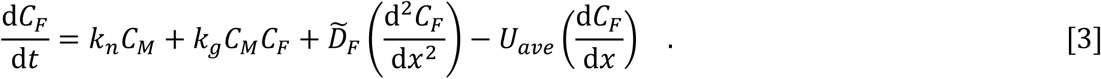

Here, *D̃_M_* and *D̃_F_* are the *effective* diffusion coefficients of the monomer and fibril, respectively, and *U_ave_* denotes the average flow velocity. The *x* shows the position measured along the axial direction of the tube. The third and fourth terms on the right-hand sides represent the effects of the self-diffusion and axial convection, respectively^55^. Aris modified the effective diffusion coefficient *D̃* proposed by Taylor^56^ and derived the following equation in a cylindrical channel with an inner diameter of β^57^:

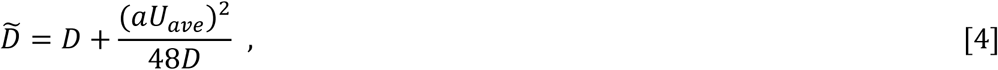

where *D* denotes the original diffusion coefficient, and this relationship was used for *D̃_M_* and *D̃_F_*.

We solved the simultaneous equation (2) and (3) numerically using the finite difference method with the model shown in Fig. 6. In the actual flow assay, the solution flows repeatedly in the looped 2.5-m tube, and the amount of the fibril is measured each time it passes through the fluorescence detector. In the numerical model, a sufficiently long (∼50 m) tube was assumed, through which monomer molecules and fibrils flowed with a carrier flow of buffer solution, causing the fibril-formation reaction, and the actual circulation behavior can be simulated by superimposing the fibril distribution every 2.5 m. As an initial condition, the monomer solution to be applied was set so that its center was 5 m from the left end of the tube, and for the boundary conditions, we assumed no flux at the ends of the tube.

## Acknowledgements

The authors would like to thank Prof. Hironobu Naiki (Fukui Univ.) and Shingi Nagamatsu (Daicel Co.) for discussion and Jun Mizunoe (Keyence Co.) for introducing fluorescence microscope. This study was performed as part of the Cooperative Research Program for the Institute for Protein Research, Osaka University, Osaka University (CR-24-02), and was supported by the Japan Society for the Promotion of Science (22H02584 to Y.Gs., 24K11427 to S.Y. and 20K06580 to K.Y.), and Daicel Co. EM measurements were carried out by using a facility in the Research Center for Ultra-High Voltage Electron Microscopy, Osaka University.

## Author contributions

Y. G., T. O., and H. O. wrote the manuscript. Y. G., T. O., I. Y. and K. Y. conducted the experiments. T. O., W. Y. and H. O. performed simulations. Y. G., K. Y., H. M., S. Y. and H. O. supervised the study.

---

**Extended Data Fig. 1.**
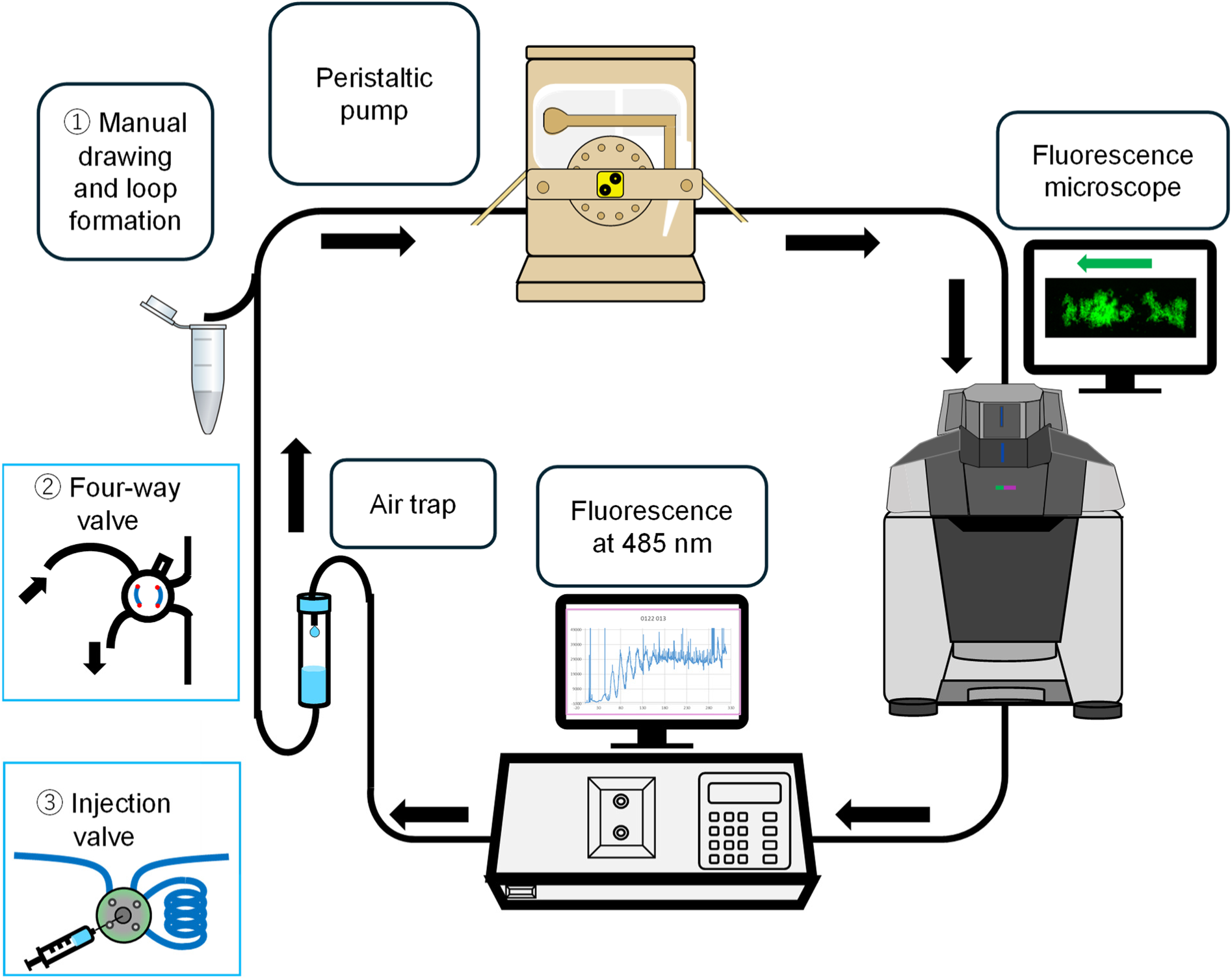
Overview of peristaltic pump-dependent amyloid inducer with the looped flow of amyloidogenic proteins. We used an ISMATEC peristaltic pump and a JASCO FP920 fluorometer with a 16 μl-volume flow cell of as the standard conditions. We also installed a Keyence fluorescence microscope to simultaneously monitor the morphology of amyloid fibrils. These apparatuses were connected by a silicone tube with an inner diameter of 1.0 nm. The total length of the loop was approximately 1.5 m with a total volume of 2-3 ml. For sample injection, we used the manual injection (1) or four-way valve (2), which could be replaced with a sample injector (3) in a future system.

**Extended Data Fig. 2.**
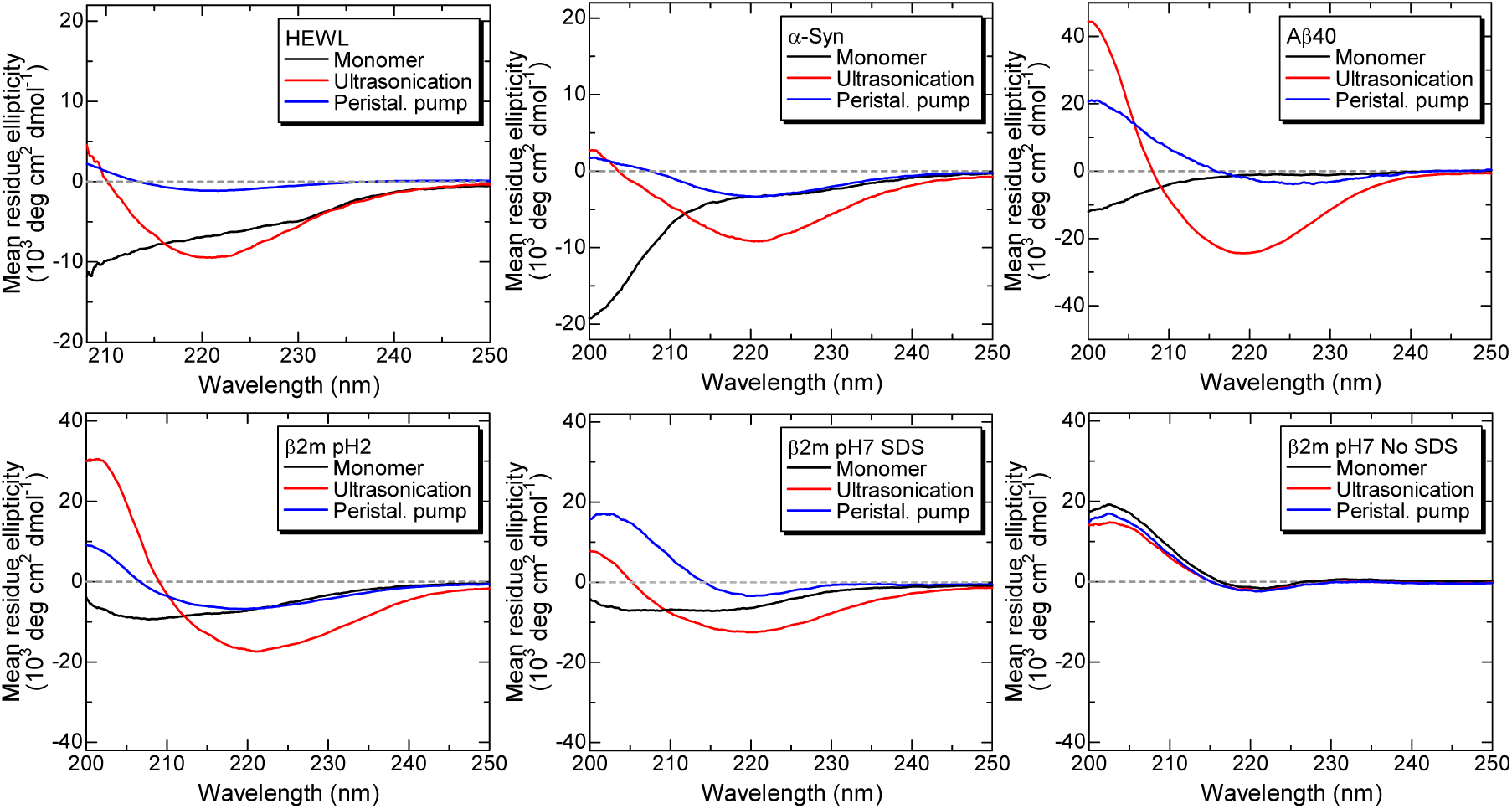
Far-UV CD spectra before and after the peristaltic pump-treatments or after ultrasonication for (a) HEWL, (b) αSN, (c) Aβ40, and (d-f) β2m under three conditions: (d) in 20 mM HCl, and (e) in the presence or (f) absence of 0.5 mM SDS at pH 7.0.

**Extended Data Fig. 3.**
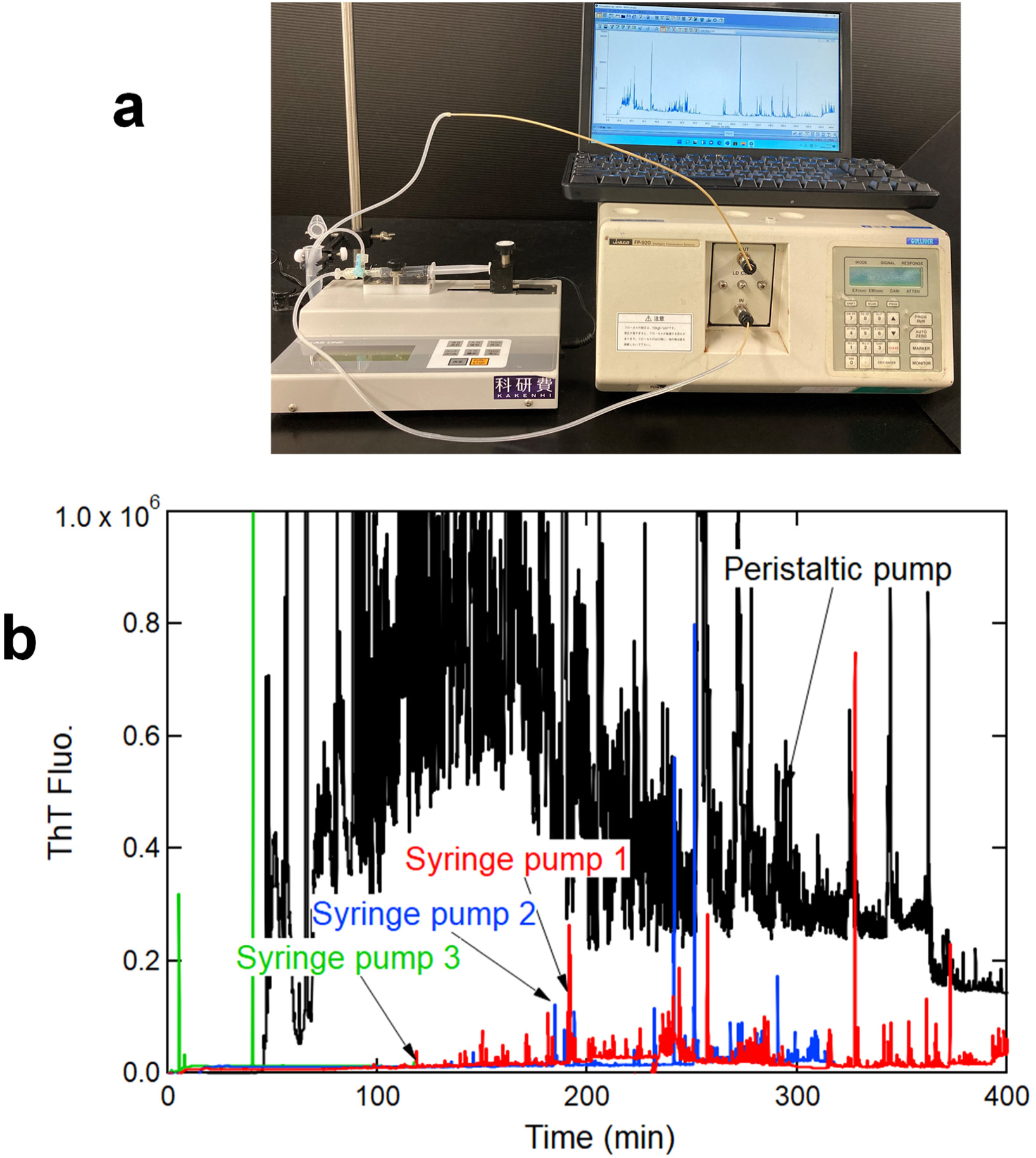
Comparison of the effects of repeated flows of the HEWL solution with syringe and peristaltic pumps. (a) A set-up photo of the syringe pump system. Totals of 2.2 ml of 1.0 mg/ml HEWL in 20 mM HCL and 2.0 M GuHCl were repeatedly flowed at 0.1 ml/min. (b) ThT fluorescence kinetics monitored at 485 nm. For the experiment with the peristaltic pump, the same HEWL solution was applied for 10 min at a flow rate of 0.1 ml/min. The total recovered volume was 2.3 ml. For the syringe pump experiments, three trials are shown with different colors.

**Extended Data Fig. 4.**
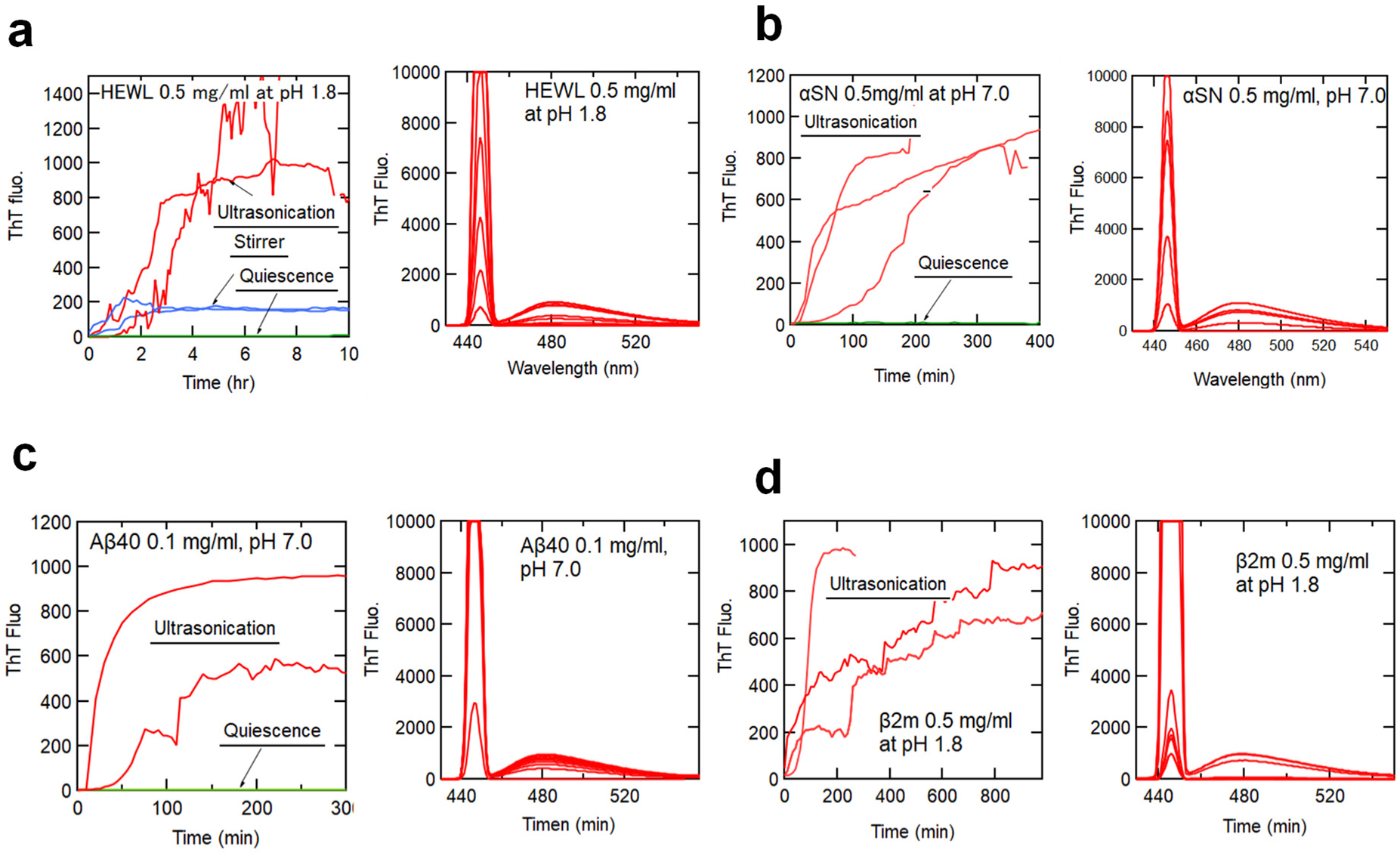
Ultrasonication-dependent amyloid formation of HEWL (a), αSN (b), Aβ40 (c), and β2m at pH 1.8 (20 mM HCl and 0.4 M NaCl) (d). Kinetics monitored by ThT fluorescence at 485 nm and spectral changes including light scattering at an excitation wavelength (445 nm) are shown. Temperature was set at 25 °C and it increased by 3-4 degrees during ultrasonication.

**Extended Data Fig. 5.**
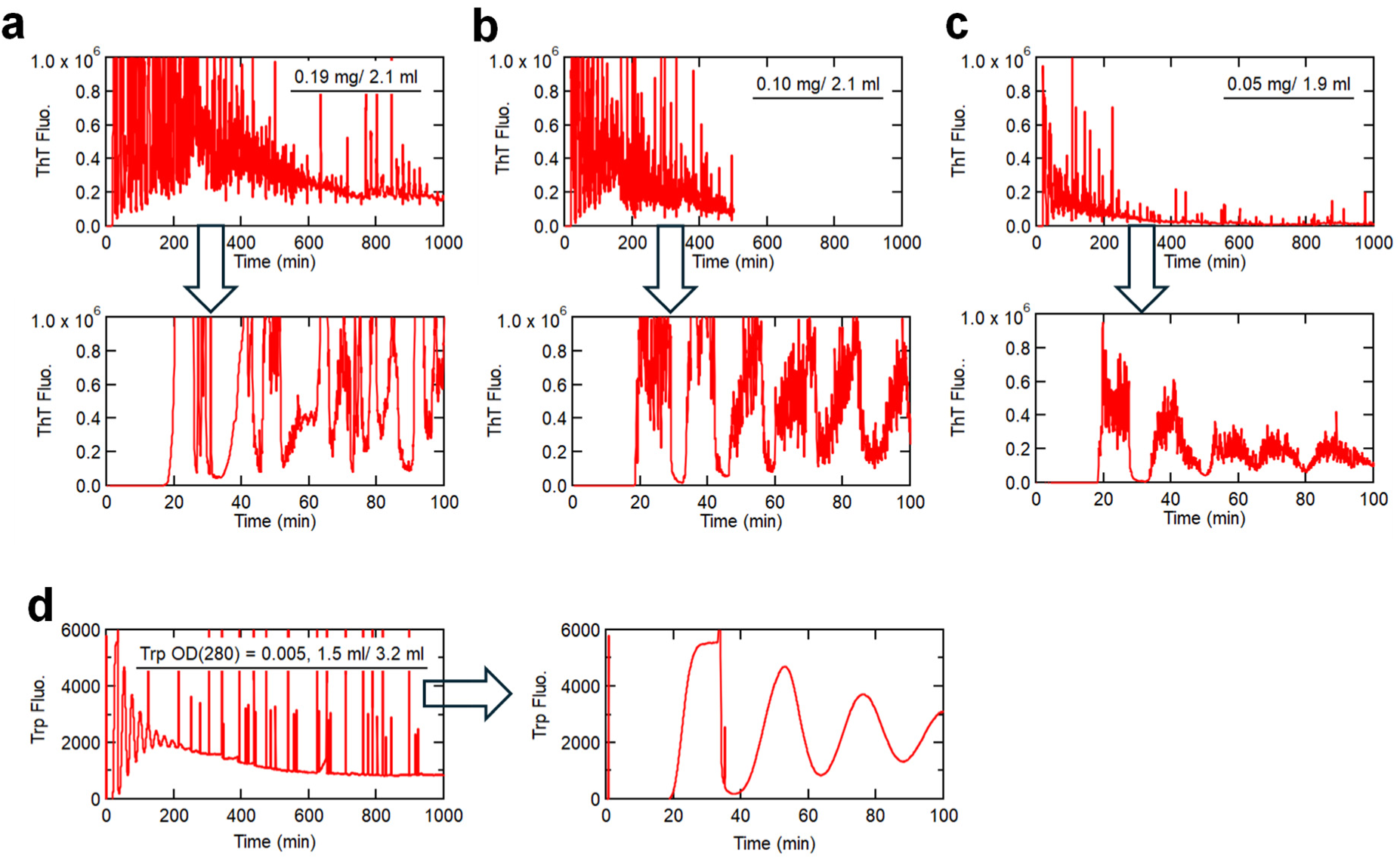
(a-c) Absorption of the preformed HEWL amyloid fibrils to the looped system at three different amounts applied. Amounts of amyloid fibrils were 0.19 mg (a), 0.10 mg (b), and 0.05 mg (c) with a loop volume of 2.0 ml. Expanded profiles at early time-periods are shown, indicating the repeated cycles of amyloid appearance with declining intensities. (d) As a control, the flow of tryptophan was monitored by its fluorescence at 350 nm.

**Extended Data Fig. 6.**
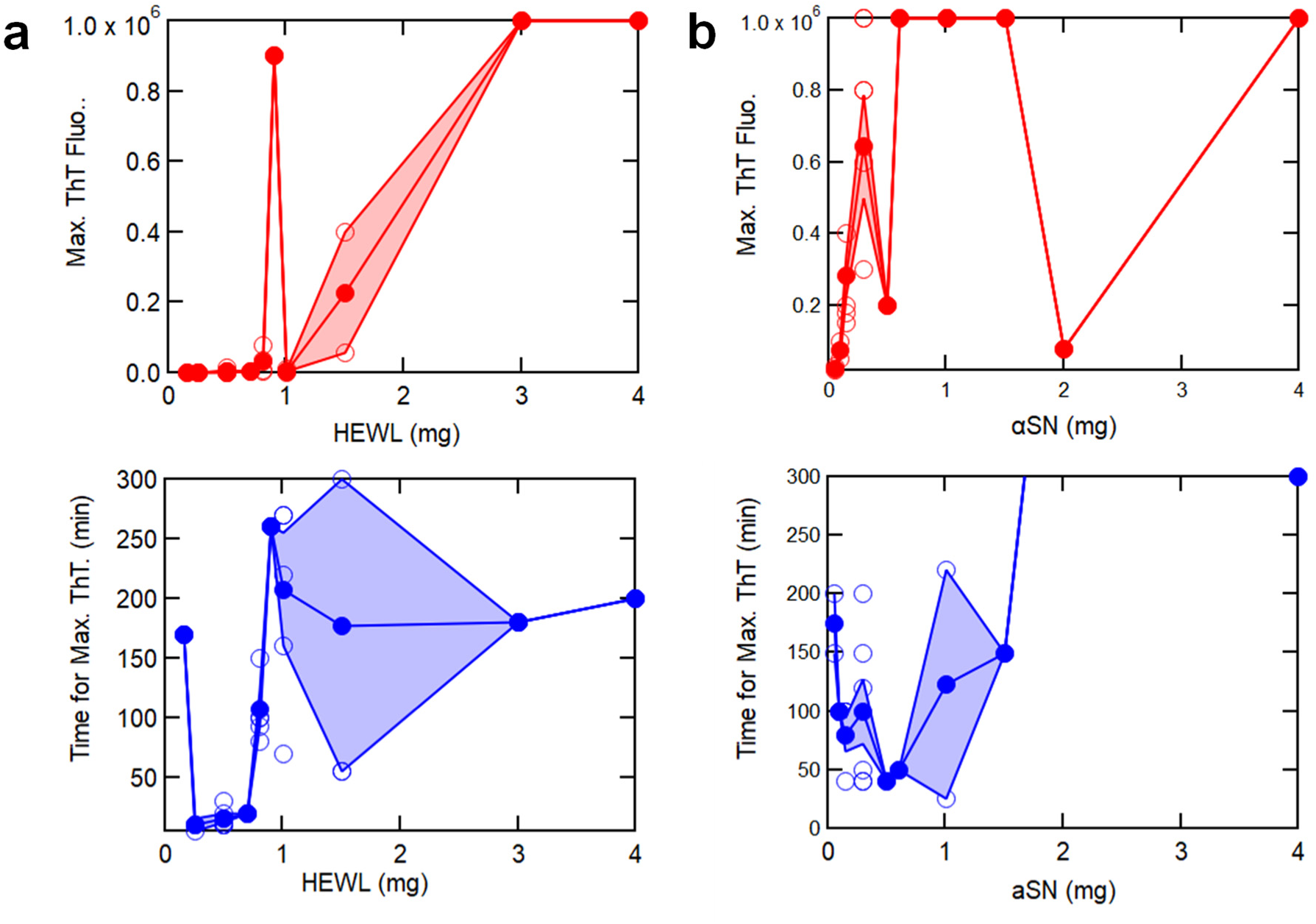
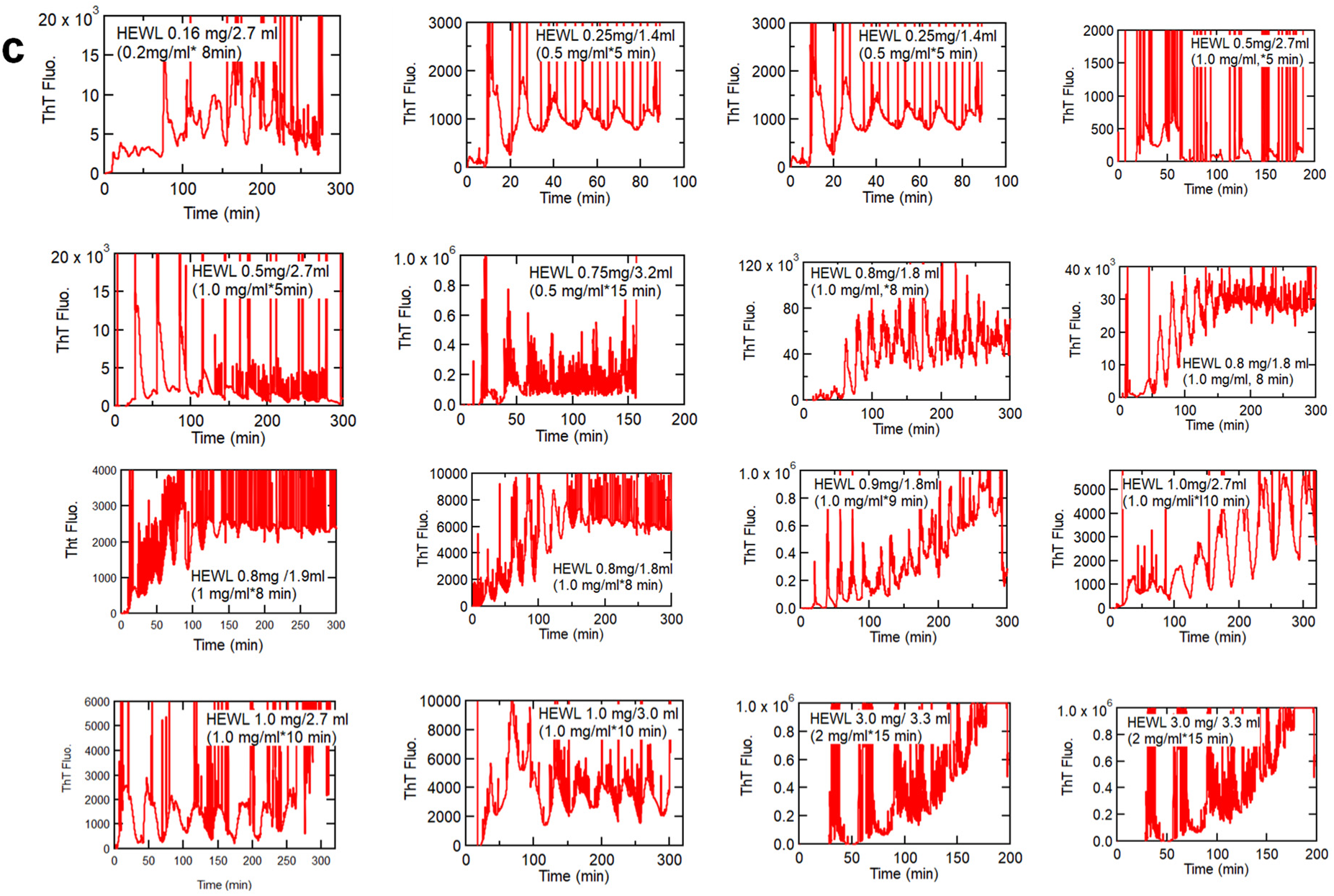
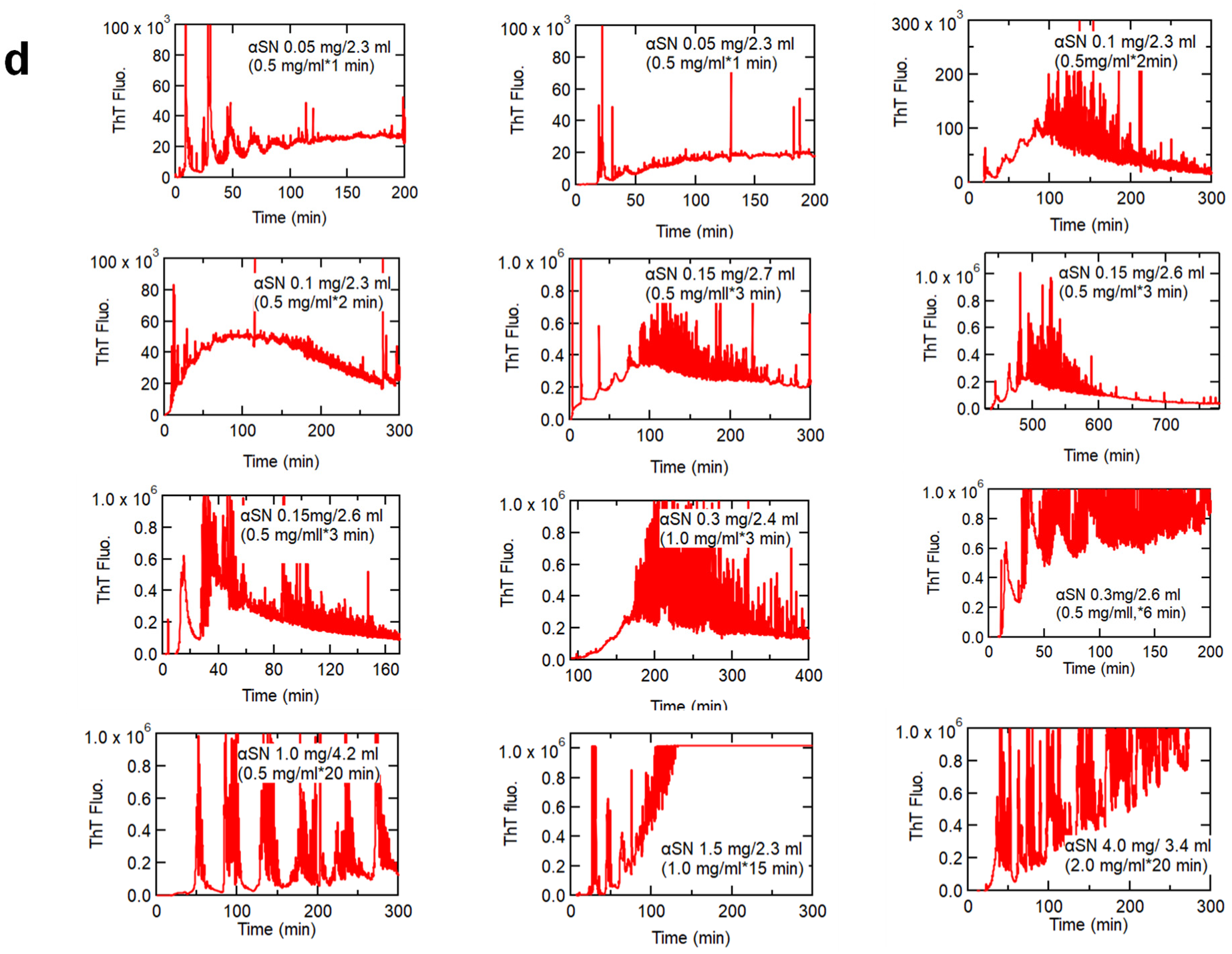
Dependencies of maximal ThT fluorescence (upper panels) and its time (lower panels) on the amount of proteins for (a) HEWL and (b) αSN. Panels (c) and (d) show some of the original data for HEWL and αSN, respectively. The kinetics at higher HEWL or αSN amounts were similar, showing the stepwise increase in peak intensity. The maximal ThT fluorescence intensity was roughly in proportion to the amount of HEWL applied. On the other hand, at lower amounts of HEWL or αSN, the first or second peak often exhibited the maximal ThT fluorescence intensity. The complicated dependence was likely to be caused by a combined effect of the intense triggering of amyloid nucleation and the absorption of amyloid fibrils to the loop surfaces.

**Extended Data Fig. 7.**
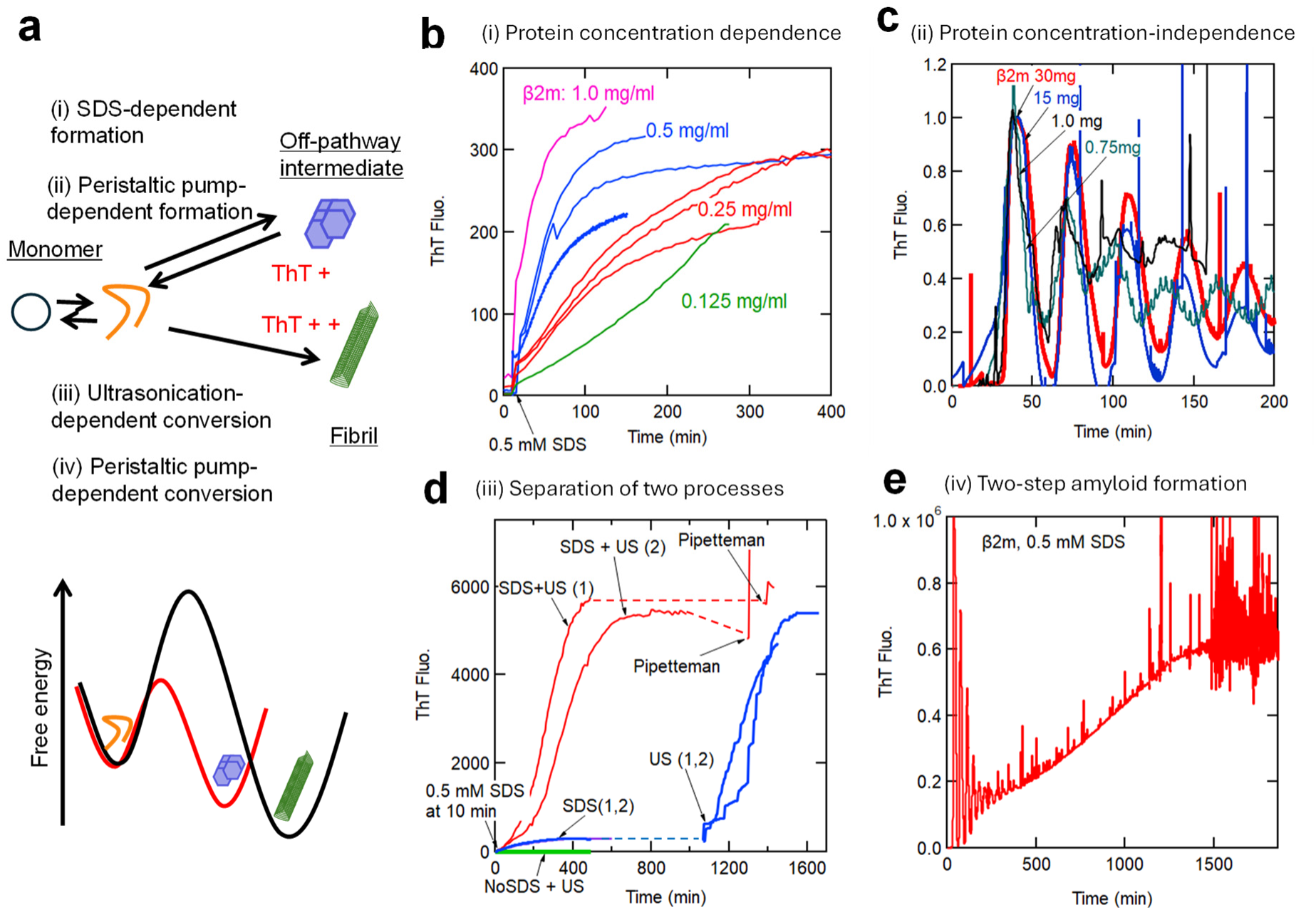
Two-step amyloid formation of β2m at pH 7.0 with an off-pathway intermediate. (a) Schematic model and a free energy diagram with an octameric off-pathway intermediate. (b) β2m concentration-dependent off-pathway intermediate formation under stirring in 0.5 mM SDS. (c) β2m concentration -independent off-pathway intermediate formation induced by the peristaltic pump. (d) Linkage and separation of ultrasonication-independent off-pathway intermediate and ultrasonication-dependent amyloid formation in 0.5 mM SDS. (e) Peristaltic pump-dependent off-pathway intermediate formation and subsequent conversion to amyloid fibrils.

**Sup. Table 1.** Comparison of amyloid fibrils prepared by the pelistaltic pump and ultrasonication. For peristaltic pump-induced amyloid fibrils, specific ThT fluorescence intensity was estimated for each recovered solution with the Hitachi F7000 fluorometer. For amyloid fibrils prepared by ultrasonication with the fluorometer, the observed values were used.

| Protein | Conditions | Methods | ThT max/(mg/mL protein) | SD | No. points |
| --- | --- | --- | --- | --- | --- |
| HEWL | 20 mM HCl (pH 2), 2.0 M Gdn-HCl | Peristaltic pump | 89 | 160 | 14 |
|  |  | Ultrasonication | 1403 | 566 | 3 |
| αSN | 50mM NaPhos. (pH 7.0), 0.5 M Na<sub>2</sub>SO<sub>4</sub> | Peristaltic pump | 1070 | 828 | 5 |
|  |  | Ultrasonication | 3692 | 2185 | 5 |
| Aβ | 20 mM NaPhos. (pH 7.0), 50 mM NaCl | Peristaltic pump | 1460 | 941 | 4 |
|  |  | Ultrasonication | 11016 | 0 | 1 |
| β2m | 20 mM HCl (pH 2), 400 mM NaCl | Peristaltic pump | 529 | 310 | 3 |
|  |  | Ultrasonication | 7485 | 2047 | 3 |
|  | 50 mM NaPhos. (pH 7.0), 100 mM NaCl, 0.5 mM SDS | Peristaltic pump | 466 | 11 | 2 |
|  |  | Ultrasonication | 16748 | 1440 | 2 |
|  | 50 mM Na Phos. (pH7), 100 mM NaCl | Peristaltic pump | 5 | 1 | 4 |
|  |  | Ultrasonication | 8 | 4 | 3 |

**Sup. Table 2.** List for experiments of peristaltic pump-dependen amyloid formation of HEWL and αSN.

|  | HEWL<br>(mg) | ThT(Max) | Max<br>time<br>(min) | (Conc. (mg/ml) * Flow<br>rate (ml/min) * Flow<br>time (min))/(Total<br>volume (ml)) | FP920<br>File No. | Notes |
| --- | --- | --- | --- | --- | --- | --- |
| 1 | 0.2 | 1,300 | 170 | 0.2*01*8 /2.71 | 118-7 |  |
| 2 | 0.3 | 1,500 | 10 | 0.5*01*5 /1.38 | 108-6 |  |
| 3 | 0.3 | 1,500 | 10 | 0.5*01*5 /1.38 | 108-8 |  |
| 4 | 0.5 | 1,600 | 10 | 1.0*01*5 /1.28 | 108-12 |  |
| 5 | 0.5 | 3,000 | 10 | 1.0*01*5 /1.35 | 108-13 |  |
| 6 | 0.5 | 1,600 | 10 | 1.0*01*5 /1.34 | 108-14 |  |
| 7 | 0.5 | 13,000 | 30 | 1.0*01*5 /2.69 | 122-8 |  |
| 8 | 0.5 | 1,800 | 20 | 1.0*01*5 /1.78 | 122-9 |  |
| 9 | 0.5 | 800 | 15 | 1.0*01*5 /1.84 | 122-11 |  |
| 10 | 0.7 | 1,900 | 20 | 1.0*01*7 /2.71 | 122-3 |  |
| 11 | 0.8 | 35,000 | 100 | 1.0*01*8 /1.82 | 122-13 |  |
| 12 | 0.8 | 78,000 | 100 | 1.0*01*8 /1.75 | 122-14 |  |
| 13 | 0.8 | 3,000 | 80 | 1.0*01*8 /1.84 | 122-15 |  |
| 14 | 0.8 | 7,000 | 150 | 1.0*01*8 /1.77 | 122-16 |  |
| 15 | 0.9 | 900,000 | 260 | 1.0*01*9 /1.79 | 122-12 |  |
| 16 | 1.0 | 10,000 | 70 | 1.0*01*10 /2.97 | 122-17 |  |
| 17 | 1.0 | 5,000 | 270 | 1.0*01*10 /2.67 | 129-2 |  |
| 18 | 1.0 | 2,000 | 270 | 1.0*01*10 /2.67 | 129-3 |  |
| 19 | 1.0 | 1,500 | 220 | 1.0*02*5 /3.05 | 129-4 |  |
| 20 | 1.4 | 650 | 13 | 1.0*0.2*7/2.59 | 206-4 |  |
| 21 | 1.5 | 55,000 | 55 | 1.0*01*15 /2.68 | 122-4 |  |
| 22 | 1.5 | 400,000 | 300 | 1.0*0.2*7.5 /3.08 | 129-5 |  |
| 23 | 3.0 | 1,000,000 | 180 | 2.0*0.1*15 /3.32 | 213-2 | Fluo. microscopy-3 |
| 24 | 4.0 | 1,000,000 | 200 | 2.0*0.2*10 /3.75 | 129-8 | Fluo. microscopy-1 |
| 25 | 4.0 | 1,000,000 | 70 | 2.0*0.2*10 /2.50 | 206-6 | Fluo. microscopy-2 |
| 26 | 7.5 | 180,000 | 40 | 0.5*0.1*15 /2.56 | 402-2 |  |
| 27 | 8.5 | 990,000 | 200 | 0.5*0.1*17 /2.06 | 402-3 |  |

| | αSN<br>(mg) | ThT(Max) | Max<br>time<br>(min) | (Conc. (mg/ml) * Flow<br>rate (ml/min) * Flow<br>time (min))/(Total<br>volume (ml)) | File No. | Notes |
| --- | --- | --- | --- | --- | --- | --- |
| 0 | 0.1 | 30,000 | 200 | 0.5*0.1*1 /2.29 | 411-4 |  |
| 1 | 0.1 | 20,000 | 150 | 0.5*0.1*1 /2.33 | 411-5-1 |  |
| 2 | 0.1 | 50,000 | 100 | 0.5*0.1*2 /2.29 | 411-2 |  |
| 3 | 0.1 | 100,000 | 100 | 0.5*0.1*2 /2.29 | 411-3 |  |
| 4 | 0.2 | 200,000 | 100 | 0.5*0.1*3 /2.41 | 407-7-1 |  |
| 5 | 0.2 | 400,000 | 100 | 0.5*0.1*3 /2.41 | 407-2 |  |
| 6 | 0.2 | 180,000 | 40 | 0.5*0.1*3 /2.41 | 407-8 |  |
| 7 | 0.2 | 150,000 | 80 | 0.5*0.1*3 /2.30 | 411-6 |  |
| 8 | 0.3 | 300,000 | 120 | 1.0*0.1*3 /2.37 | 407-3-1 |  |
| 9 | 0.3 | 1,000,000 | 150 | 1*0.1*3 /2.21 | 407-5-1 |  |
| 10 | 0.3 | 800,000 | 200 | 1*0.1*3 /2.21 | 407-5-2 |  |
| 11 | 0.3 | 800,000 | 40 | 0.5*0.1*6 /2.64 | 407-6-1 |  |
| 12 | 0.3 | 1,000,000 | 50 | 0.5*0.1*6 /2.64 | 407-6-2 |  |
| 13 | 0.3 | 600,000 | 40 | 0.5*0.1*6 /2.30 | 411-6-2 |  |
| 14 | 0.5 | 200,000 | 40 | 0.5*0.1*10 /4.3 | 229-4 |  |
| 15 | 0.6 | 1,000,000 | 50 | 0.5*0.1*12 /2.32 | 411-5-2 |  |
| 16 | 1.0 | 1,000,000 | 50 | 0.5*0.1*20 /4.19 | 221-2 | Fluo. microscopy-3 |
| 17 | 1.0 | 1,000,000 | 220 | 0.5*0.1*20 /4.52 | 227-2 | Fluo. microscopy-4 |
| 18 | 1.0 | 1,000,000 | 25 | 0.5*0.1*20 /4.08 | 229-2 | Fluo. microscopy-5 |
| 19 | 1.5 | 1,000,000 | 150 | 1*0.1*15 /2.27 | 402-4 |  |
| 20 | 2.0 | 80,000 | 600 | 2*0.1*10 /3.21 | 215-4 | Fluo. microscopy-1 |
| 21 | 4.0 | 1,000,000 | 300 | 2*0.1*20 /3.42 | 219-2 | Fluo. microscopy-2 |

| | Aβ<br>(mg) | ThT(Max) | Max<br>time<br>(min) | (Conc. (mg/ml) * Flow<br>rate (ml/min) * Flow<br>time (min))/(Total<br>volume (ml)) | File No. | Comments |
| --- | --- | --- | --- | --- | --- | --- |
| 1 | 0.1 | 840,000 | 55 | 0.086*0.1*10/2.5 | 328-2 | DMSO for dissolution |
| 2 | 0.1 | 1,000,000 | 20 | 0.086*0.1*10/2.5 | 329-2 | DMSO for dissolution |
| 3 | 0.1 | 1,000,000 | 25 | 0.086*0.1*10/2.5 | 401-2 | DMSO for dissolution |
| 4 | 0.1 | 370,000 | 71 | 0.086*0.1*10/3 | 418-3 | Fluo. microscopy-1 |
| 5 | 0.1 | 1,000,000 | 26 | 0.086*0.1*10/2.5 | 709-2 |  |
| 6 | 0.1 | 700,000 | 23 | 0.086*0.1*10/2.44 | 710-3 |  |
| 7 | 0.1 | 1,000,000 | 25 | 0.086*0.1*10/2.23 | 710-4 |  |
| 8 | 0.1 | 5,000,000 | 60 | 0.09*0.1*15/2.8 | 725-2 | Fluo. microscopy-2 |
| 9 | 0.2 | 1,000,000 | 45 | 0.13*0.1*15/2.85 | 726-2 | Fluo. microscopy-3 |

| | β2m (mg) | ThT(Max) | Max time (min) | (Conc. (mg/ml) * Flow rate (ml/min) * Flow time (min))/(Total volume (ml)) | File No. | Comments |
| --- | --- | --- | --- | --- | --- | --- |
|  | pH 1.8 |  |  |  |  |  |
| -2 | 0.5 | 300,000 | 150 | 0.5*0.1*10/2.09 | 202-3 |  |
| -1 | 0.5 | 400,000 | 150 | 0.5*0.1*10/1.78 | 202-5 |  |
| 0 | 0.5 | 1,000,000 | 105 | 0.5*0.1*10/2.04 | 520-2 |  |
| 1 | 0.5 | 760,000 | 107 | 0.5*0.1*10/2.02 | 521-2 |  |
| 2 | 0.5 | 650,000 | 217 | 0.5*0.1*10/2.97 | 529-2 | Fluo. microscopy |
| 3 | 0.8 | 800,000 | 100 | 0.5*0.1*15/2.17 | 202-4 |  |
|  | pH 7.0, 0.5 mM SDS |  |  |  |  |  |
| 1 | 0.5 | 4,000 | 22 | 0.5*0.1*10/2.70 | 206-8 |  |
| 2 | 0.5 | 1,500 | 22 | 0.5*0.1*5/2.67 | 206-9 |  |
| 3 | 1.0 | 150,000 | 40 | 0.5*0.1*20/3.59 | 304-2 |  |
| 4 | 15.0 | 1,000,000 | 640 | 10*0.1*15/2.67 | 516 | Fluo. microscopy-2 |
| 5 | 30.0 | 1,000,000 | 1800 | 20*0.1*15/2.82 | 530 | Fluo. microscopy-1 |
|  | pH 7.0, No SDS |  |  |  |  |  |
| -1 | 0.5 | 2,000 | 15 | 0.5*0.1*9 | 206-10 |  |
| 0 | 0.8 | 9,700 | 34 | 0.5*0.1*15/2.4 | 826-3 |  |
| 1 | 0.8 | 9,400 | 32 | 0.5*0.1*15/2.5 | 826-4 |  |
| 2 | 1.0 | 5,000 | 33 | 0.5*0.1*20/3.19 | 215-2 |  |
| 3 | 3.0 | 5,000 | 35 | 2.0*0.1*15/2.95 | 215-3 |  |
| 4 | 3.0 | 6,700 | 30 | 2.0*0.1*15/1.74 | 427-4 |  |
| 5 | 15.0 | 88,000 | 23 | 10*0.1*15/1.8 | 430-3 |  |
| 6 | 15.0 | 70,000 | 50 | 10*0.1*15/2.95 | 501 | Fluo. microscopy-1 |
| 7 | 15.0 | 100,000 | 35 | 10*0.1*15/2.7 | 516-2 |  |
| 8 | 30.0 | 150,000 | 40 | 20*0.1*15/3.16 | 502 | Fluo. microscopy-2 |

